# Refining the pool of RNA-binding domains advances the classification and prediction of RNA-binding proteins

**DOI:** 10.1101/2023.08.17.553134

**Authors:** Elsa Wassmer, Gergely Koppàny, Malte Hermes, Sven Diederichs, Maïwen Caudron-Herger

## Abstract

**Key Points:**

- Comprehensive analysis of RNA-related protein domains and families enriched in RNA-binding proteins (RBPs)
- Pan-species prediction of new RBPs, and prediction and validation of new RNA-binding domains
- Online resource with complete dataset including high-confidence human RBPs according to a new scoring system

From transcription to decay, RNA-binding proteins (RBPs) influence RNA metabolism. Using the RBP2GO database that combines proteome-wide RBP screens from 13 species, we investigated the RNA-binding features of 176896 proteins. By compiling published lists of RNA-binding domains (RBDs) and RNA-related protein family (Rfam) IDs with lists from the InterPro database, we analyzed the distribution of the RBDs and Rfam IDs in RBPs and non-RBPs to select RBDs and Rfam IDs that were enriched in RBPs. We also explored proteins for their content in intrinsically disordered regions (IDRs) and low complexity regions (LCRs). We found a strong positive correlation between IDRs and RBDs and a co-occurrence of specific LCRs. Our bioinformatic analysis indicated that RBDs/Rfam IDs were strong indicators of the RNA-binding potential of proteins and helped predicting new RBP candidates, especially in less investigated species. By further analyzing RBPs without RBD, we predicted new RBDs that were validated by RNA-bound peptides. Finally, we created the RBP2GO composite score by combining the RBP2GO score with new quality factors linked to RBDs and Rfam IDs. Based on the RBP2GO composite score, we compiled a list of 2018 high-confidence human RBPs. The knowledge collected here was integrated into the RBP2GO database at https://RBP2GO-2-Beta.dkfz.de.

GRAPHICAL ABSTRACT

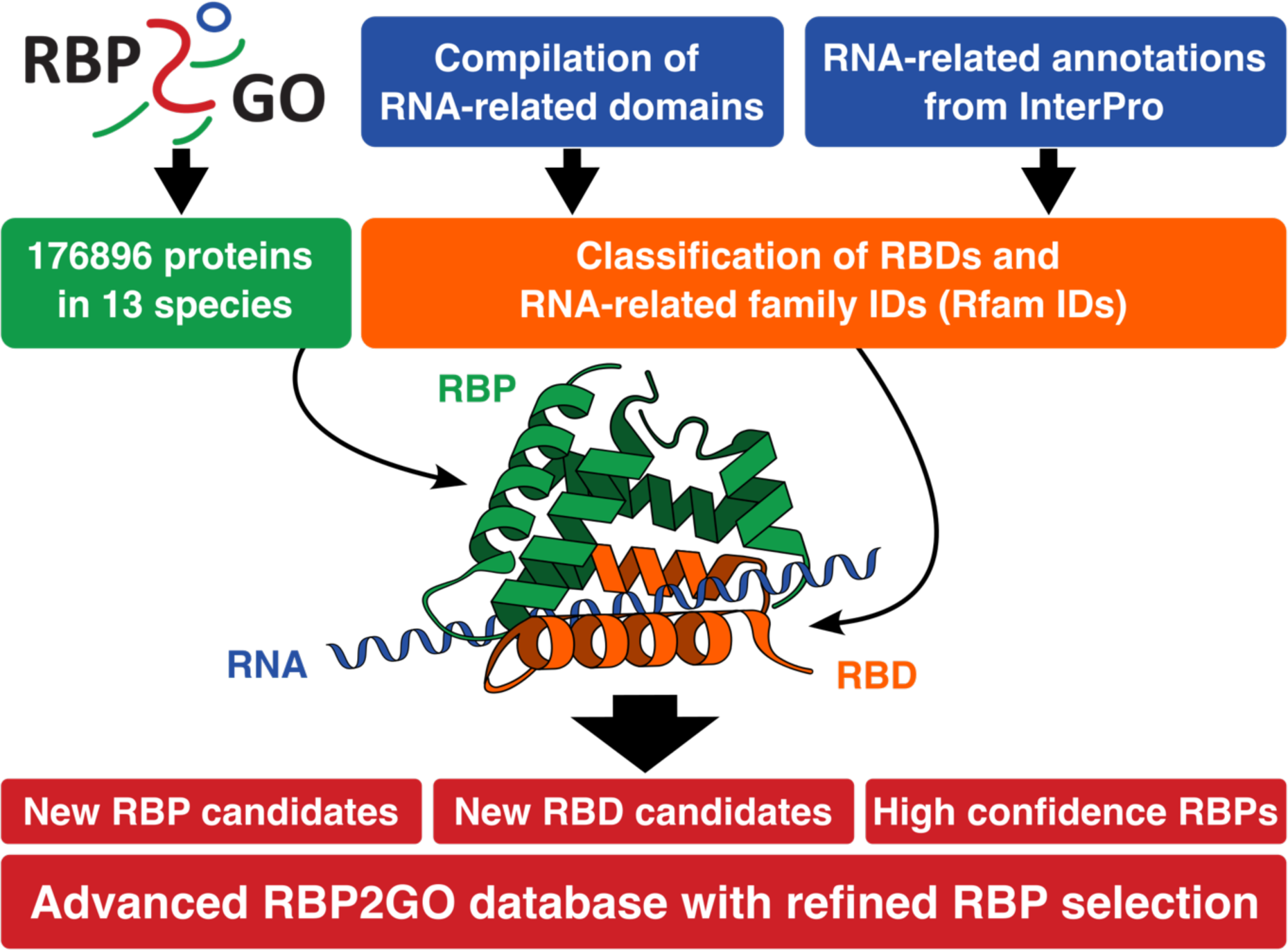

## INTRODUCTION

RNA and RNA-binding proteins (RBPs) have emerged as central players in a large number of key cellular processes (1) with significant implications for our understanding of severe diseases such as neurological disorders or cancer (2–6). Therefore, it is fundamental to comprehensively examine RNA-protein complexes and their molecular functions.

Over the last decade, a variety of orthogonal techniques were developed to systematically identify RBPs based on proteome-wide analyses. The study of the mRNA-bound proteome (7–12) combines the UV-crosslinking of RNA molecules and their bound proteins with the use of oligo-dT beads to isolate the RNA-protein complexes and identify the bound proteins via mass spectrometry. The technique was further improved to identify the RNA-binding regions within the RBPs (13) and was applied on different human cells such as HEK293, HeLa and K562 (7,8,12), as well as other model organisms such as *Mus musculus* (10,14,15). Other strategies were explored, notably the isolation of crosslinked RNA-RBP complexes by organic phase separation (16–18), or the separation of RNA-dependent complexes on sucrose gradients upon RNase treatment (19–21).

These massive efforts resulted in the generation of an ever-growing number of datasets. In particular for model species such as *Homo sapiens*, six datasets were available in 2018 (13) as compared to 43 in 2020 (22), whereas only three datasets had been published for *Danio rerio* (23). Furthermore, previous work showed the poor overlap between the different datasets available for *Homo sapiens,* with many proteins being detected only once as RBP (22), highlighting the need for tools that could quantify the relevance of RBP candidates. A number of prediction algorithms were developed, mostly based on the protein sequence (24). Alternatively, the SONAR study predicted RNA-binding properties based on protein-protein interaction, as RBPs often interact with other RBPs (19,25). To facilitate the efficient screening of the remarkably growing RBP resources, we recently compiled all the available datasets into the comprehensive pan-species RBP2GO database (https://rbp2go.dkfz.de). RBP2GO encompasses 103 datasets and 22554 RBP candidates, along with their interactions and their functions (22). The database also displays the RBP2GO score for each protein, which reflects the probability for an RBP candidate to be a true RBP (22). However, due to a lack of data for some species, this score is only available for 9 out of the 13 featured species. Furthermore, it does not take into account data on the functional domains of the proteins.

A number of canonical RNA-binding domains (RBDs) such as the RNA-recognition motif (RRM) or the K-homology (KH) domain (26,27) have been catalogued (28) and further exploited to identify new RBPs (29). However, recent studies revealed that non-canonical RBDs and intrinsically disordered regions (IDRs) were also enriched in RBP candidates (8,30). Moreover, IDR-containing proteins such as NF-kappa-B-activating protein (NKAP) can bind to RNA in the absence of RBD via their IDRs (31). The potential of IDRs for binding to RNA was further confirmed in proteome-wide studies that directly identified peptides bound to RNA in *Homo sapiens* and *Drosophila melanogaster* (13,32). Currently, many of the newly identified RBP candidates are awaiting further experimental validation. Assessing the specificity of the rapidly growing list of RBP candidates urgently needs further criteria to evaluate the likelihood of each protein to be a true RBP.

In this study, we addressed this issue by performing a pan-species analysis of the RNA-binding characteristics of the proteins included in the RBP2GO database. To this aim, we collected lists of RBDs and RNA-related protein families (Rfam) from published studies (8,13,29) and from the InterPro database (33). We examined their distribution in RBP candidates as compared to non-RBPs in order to compile a list of selected RBDs and Rfam identifiers (Rfam IDs) that were enriched in RBP candidates. Using data from the MobiDB database (34), we also analyzed the IDR and low complexity region (LCR) content and their distribution in proteins with respect to their RBD content and found a strong correlation between the IDR/LCR and RBD content. Our bioinformatic approach indicates that the presence of a selected RBD / Rfam ID can be used to evaluate the RNA-binding potential of proteins as well as to predict new RBP candidates, in particular in species with very few available proteome-wide studies. In addition, we analyzed the RBP candidates lacking selected RBDs and identified 15 RBDs that were not in the starting collection of RBDs, that we further validated based on studies of RNA-bound peptides. Finally, by calculating a new RBP2GO composite score that combines the former RBP2GO score (19) together with new quality factors characterizing the selected RBDs and Rfam IDs, we computed a list of 2018 high-confidence human RBPs. The results of our analysis were implemented into the RBP2GO database, along with new search options to browse 1) RBDs and Rfam IDs via a given InterPro ID and 2) proteins, with respect to their RNA-binding features. We anticipate that this additional information will considerably help refining our classification of RBP candidates and foster new developments in the field.

## MATERIAL AND METHODS

### Compilation of a list of RNA-binding domain candidates

Based on a literature search, the lists of RBDs from three studies were downloaded from the respective supplementary information (8,13,29). The domain IDs were converted into InterPro IDs if referenced from another database such as Pfam (35) and finally fused together with the list of IDs from InterPro. Outdated IDs were updated or discarded if no corresponding InterPro ID could be found. Altogether, 808 IDs were retrieved from the published datasets.

Based on a database search, we first downloaded the list of the InterPro IDs related to “RNA-binding” from the InterPro website (https://www.ebi.ac.uk/interpro/, InterPro v.88 released on March 10^th^, 2022) (33). Second, since the InterPro database provides a Gene Ontology (GO) annotation that is associated to the InterPro IDs, we downloaded all the IDs annotated with the GO term “RNA-binding” (GO:0003723). Third, we downloaded from the InterPro website all the domains that overlapped with the “RNA binding domain superfamily” (IPR035979). Taken together, these three datasets amounted to 2251 unique InterPro IDs.

The fusion of the IDs from the InterPro database and the published datasets resulted in a list of 2712 unique RNA-related IDs (Supplementary Table S1). These RNA-related InterPro IDs were further filtered for the “Domain” and “Repeat” types of IDs (1289 IDs, Figure 1A).

**Figure 1.**
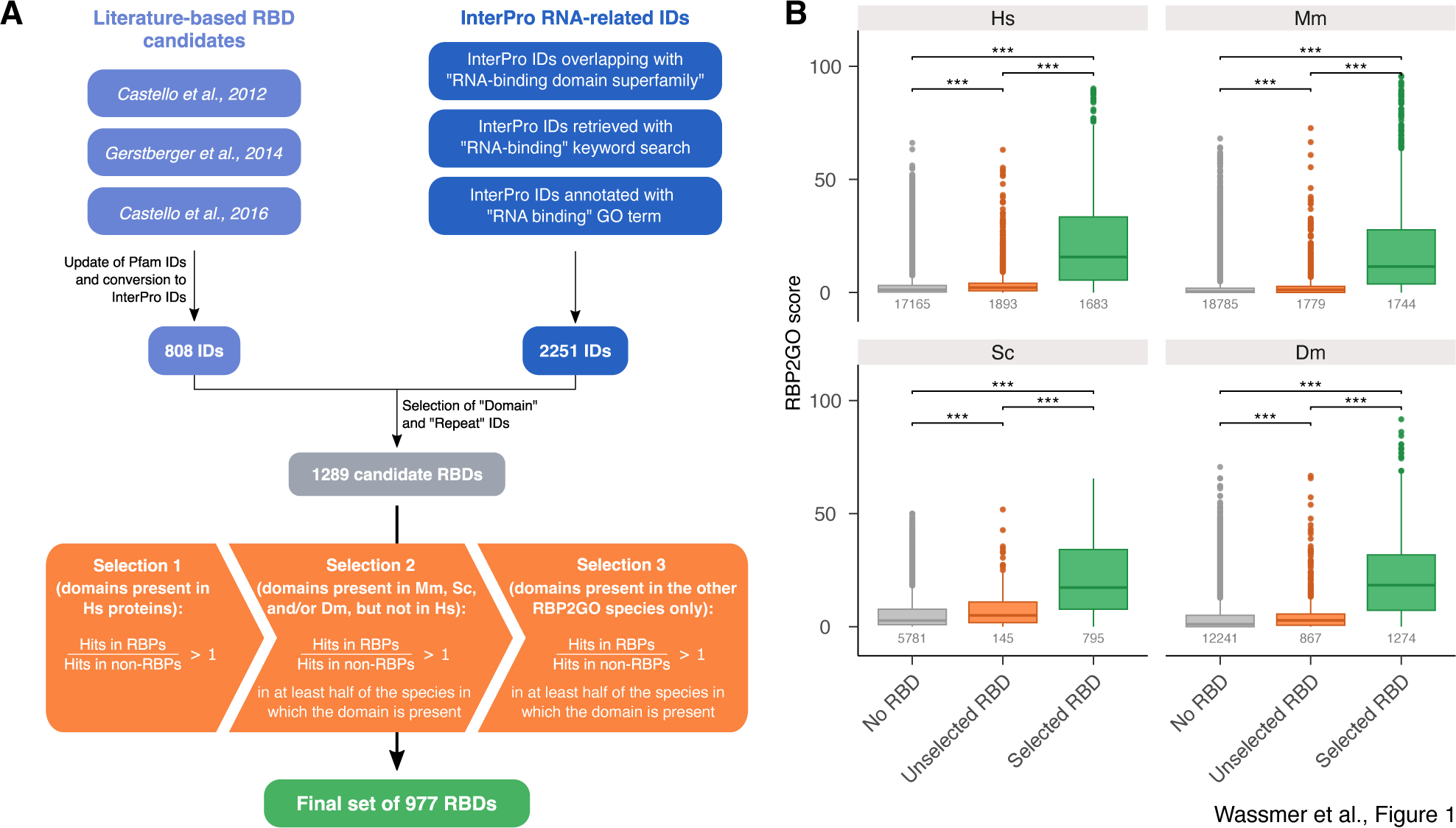
Selection of the RNA-binding domains. (**A**) Flow-chart showing the selection process of the RBD candidates and depicting the three-step selection procedure based on the ratios between RBPs and non-RBPs in the species available in the RBP2GO database (22). The starting lists of RBD candidates and the final list of selected RBDs are found in Supplementary Table S1 and Supplementary Table S3, respectively. (**B**) Boxplots depicting the distribution of the RBP2GO score of the proteins from the groups with no candidate RBDs (grey, no RBD), unselected candidate RBDs (orange) and selected RBDs (green) in Hs (*Homo sapiens*), Mm (*Mus musculus*), Sc (*Saccharomyces cerevisiae*) and Dm (*Drosophila melanogaster*). The numbers below the boxplots indicate the size of the groups. *, ** and *** correspond to *P*-values < 0.05, 0.01 and 0.001, respectively, as resulting from a Wilcoxon rank sum test.

### Selection of the RNA-binding domains

A three-step selection procedure was applied to the set of RBD candidates based on the hit ratios between RBPs (RBP candidates from proteome-wide studies, i.e., detected in at least one study) and non-RBPs in the species reported in the RBP2GO dataset (https://rbp2go.dkfz.de) (22). First, we selected the InterPro IDs of the RBD candidates that were enriched (hits in RBPs > hits in non-RBPs) in *Homo sapiens* (Hs) (Selection 1, Figure 1A) as the number of studies available for this species was by far the largest as compared to the other species (Supplementary Figure S1). We updated the UniProt IDs of the proteins according to the UniProt release 2022_01 from February 23^rd^, 2022 (36). The data for the obsolete UniProt IDs were either deleted when the ID was fused to another one already existing in the database, or duplicated if the ID was split between several new UniProt IDs. The updated sizes of the proteomes are reported in Supplementary Table S2. In a second step, for RBDs not present in Hs (Selection 2, Figure 1A), we selected the InterPro IDs of the RBD candidates that were enriched in *Mus Musculus* (Mm), *Saccharomyces cerevisiae* (Sc) and *Drosophila melanogaster* (Dm) in at least half of the species where the RBD was present. Finally, for RBDs present in neither of the four species Hs, Mm, Sc, Dm (Selection 3, Figure 1A), we selected the RBDs that were enriched in at least half of the remaining nine species where the RBD was present, *i.e.*, *Arabidopsis thaliana* (At), *Caenorhabditis elegans* (Ce), *Plasmodium falciparum* (Pf), *Escherichia coli* (Ec), *Danio rerio* (Dr), *Trypanosoma brucei* (Tb), *Salmonella Typhimurium* (ST), *Leishmania mexicana* (Lm) and *Leishmania donovani* (Ld). This three-step selection procedure led to the selection of 977 RBDs (Supplementary Table S3). The proteins of the species represented in the RBP2GO database were updated with the information on the selected RBDs (Supplementary Table S4). The data available in the Supplementary Table S4 was used to produce the graphs displayed in Figures 1B and 2A-D.

**Figure 2.**
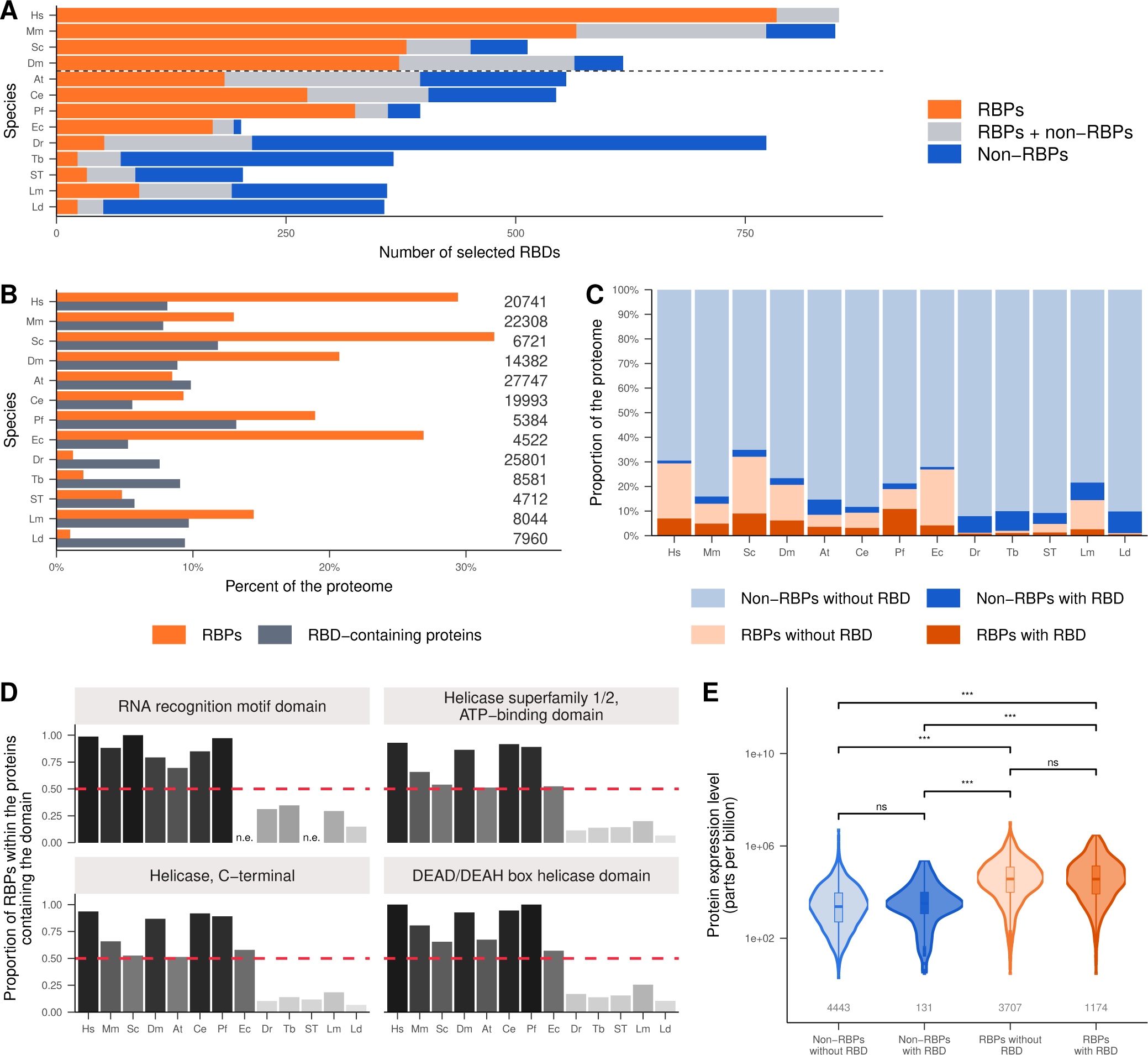
Distribution of the selected RBDs in the proteins of the RBP2GO database. (**A**) Number of selected RBDs present in each species and grouped according to the presence of the RBD either only in RBPs (orange), in both RBPs and non-RBPs (gray) or only in non-RBPs (blue). The dashed line represents the separation between the species with most (top) and fewer studies available. (**B**) Percentage of the proteome of the respective species comprised by RBPs (orange) or RBD-containing proteins (grey). The numbers at the end of the bars indicate the size of the respective proteomes in total. (**C**) Proportion in % of non-RBPs without RBD (light blue), non-RBPs with RBD (dark blue), RBPs without RBD (light orange) and RBPs with RBD (dark orange) in the different proteomes. (**D**) Fraction of RBPs among all proteins containing the indicated RBDs in the listed species. The percentages are depicted from black (high) to light grey (low). n.e. = non-existent, *i.e.,* the given domain is absent from the species. (**E**) Violin plots representing the protein expression level in HeLa cells according to a mass spectrometry experiment (37) in the four protein groups: non-RBPs without RBD (light blue), non-RBPs with RBD (dark blue), RBPs without RBD (light orange) and RBPs with RBD (dark orange). *** correspond to *P*-values < 0.001, ns corresponds to non-significant, as resulting from a Wilcoxon rank sum test.

### Protein expression levels in HeLa cells

The expression data displayed in the Figure 2E and Supplementary Figure S2 was downloaded for analysis from the EBI atlas website (https://www.ebi.ac.uk/gxa/experiments/E-PROT-19/Results) (37). The expression values can be found in the Supplementary Table S5.

### Retrieval of the InterPro domain coordinates and compilation of the number and content fraction of RBDs per protein

The coordinates of each InterPro ID of the selected RBDs in the proteins were downloaded from the InterPro database (33) (https://ftp.ebi.ac.uk/pub/databases/interpro/releases/88.0/, released on March 10^th^, 2022).

To compute the RBD content fraction per protein, we merged separately all the overlapping RBDs present in a protein. Next, we used the protein length previously retrieved from UniProt (36) (https://ftp.ebi.ac.uk/pub/databases/uniprot/previous_releases/release-2022_01/, released on February 23^rd^, 2022) to calculate the fraction of the sequence annotated as RBD for each protein.

To compute the number of RBDs present in each protein, we merged the overlapping coordinates for the InterPro ID in R using the “bed_merge()” function from the “valr” package (38). We then merged, for each protein, the coordinates of the InterPro IDs overlapping by more than 10 amino acids using the same function. The number of coordinate pairs (start, end) generated was then calculated to obtain the number of RBDs per protein which were independent in the sense that they were not overlapping by more than 10 amino acids.

The Supplementary Table S4 was updated with this information and used to produce the Figure 3 and Supplementary Figure S3.

**Figure 3.**
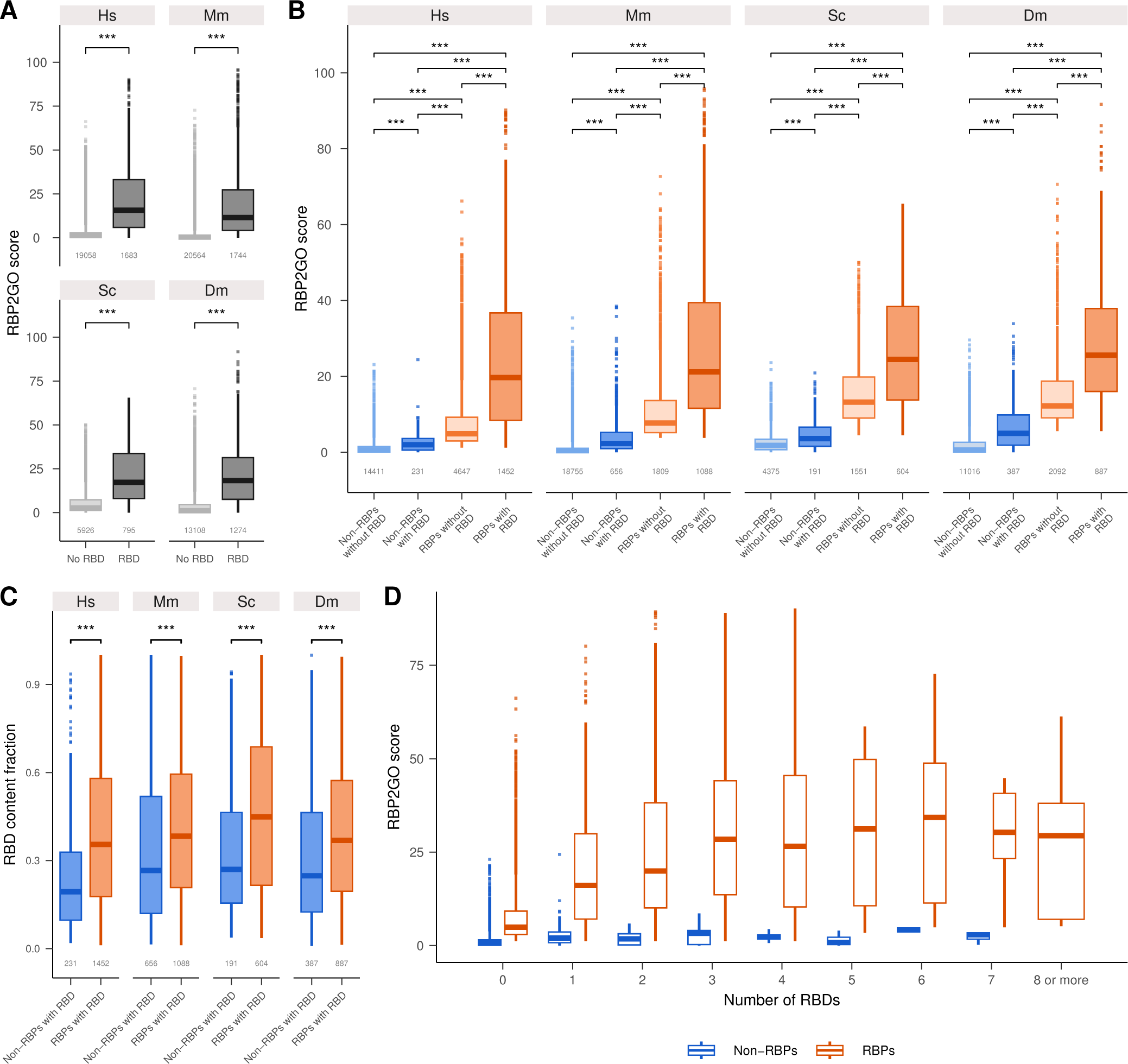
Correlation of the RBDs in the proteins of the well-studied species of the RBP2GO database with the RBP2GO score. (**A**) Boxplots representing the distribution of the RBP2GO score in proteins without RBD (light grey) and with RBD (dark grey). (**B**) Same as in (**A**), but separated into non-RBPs without RBD (light blue), non-RBPs with RBD (dark blue), RBPs without RBD (light orange) and RBPs with RBD (dark orange). (**C**) Boxplots representing the RBD content fraction per protein in non-RBPs with RBD (blue) and RBPs with RBD (orange). (**D**) Boxplots representing the distribution of the RBP2GO score per number of RBDs in non-RBPs (blue) and in RBPs (orange) in *Homo sapiens* (HS). *** corresponds to *P*-values < 0.001 as resulting from a Wilcoxon rank sum test.

### Compilation of a list of RNA-related family IDs and selection of the IDs enriched in RBP candidates

In addition to the domains and repeats InterPro ID types (see above), we computed a list of RNA-related InterPro family IDs or Rfam IDs. The “RNA binding domain superfamily” (IPR035979) overlaps with several other domains. Accordingly, the list of 2712 RNA-related InterPro IDs (Supplementary Table S1) was filtered to keep only 1028 family IDs. The same procedure as the one used for the selection of the RBDs was applied to select for the Rfam IDs that were enriched in RBP candidates. For more simplicity in the text, RBP candidates (including RBPs and non-validated RBPs) will be referred to as RBPs. This resulted in a final list of 672 Rfam IDs (Supplementary Table S6). The Supplementary Table S4 was updated accordingly with this information.

The results of these analyses were used to produce Figure 4 and Supplementary Figures S4 and S5.

**Figure 4.**
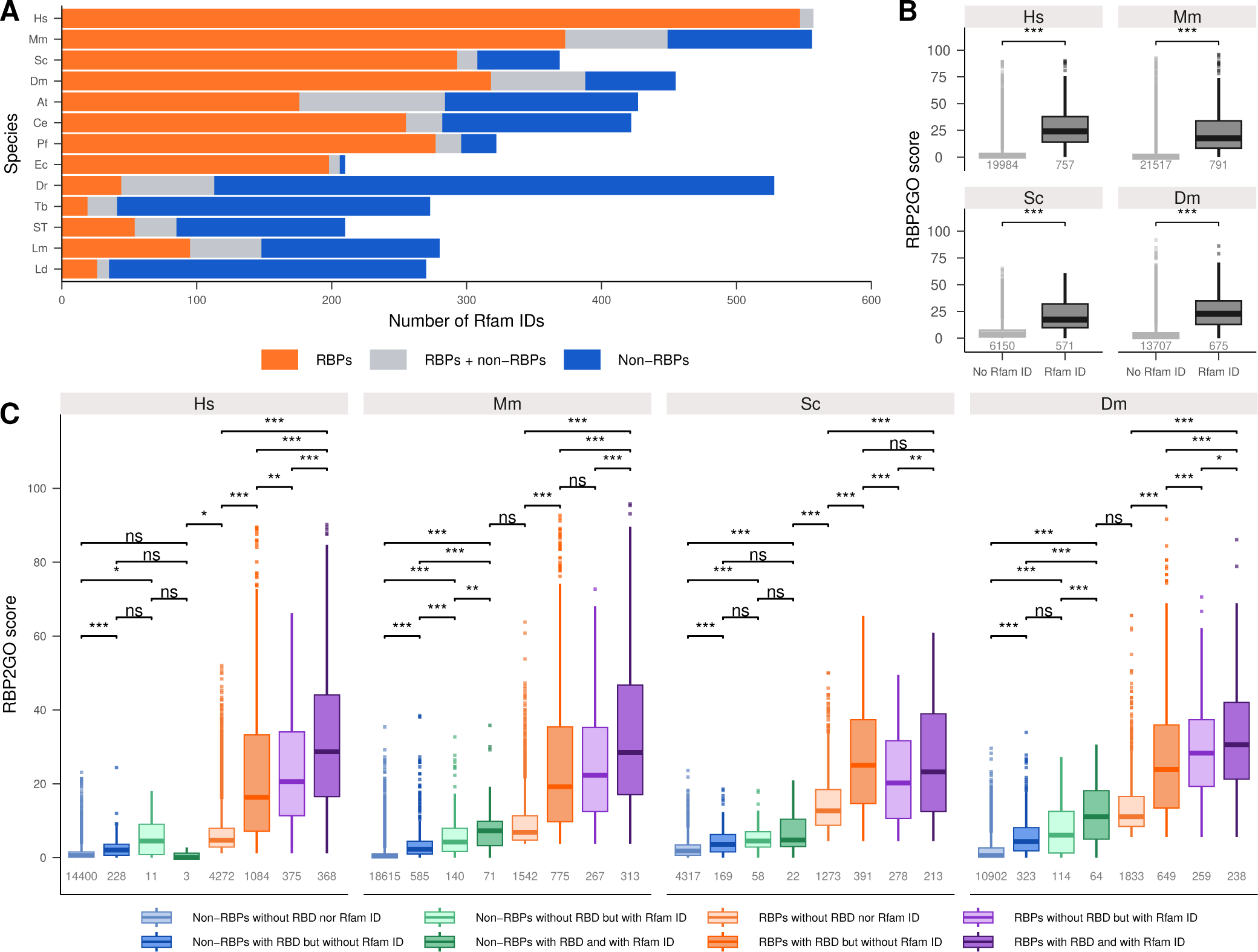
Distribution of the selected Rfam IDs in the proteins of the RBP2GO database. (**A**) Number of selected Rfam IDs present in each species and grouped according to the presence of the Rfam ID either only in RBPs (orange), in both RBPs and non-RBPs (gray) or only in non-RBPs (blue). (**B**) Boxplots representing the distribution of the RBP2GO score in proteins without Rfam ID (light grey) and with Rfam ID (dark grey). (**C**) Boxplots representing the distribution of the RBP2GO score in non-RBPs without RBD nor Rfam ID (light blue), non-RBPs with RBD but without Rfam ID (dark blue), non-RBPs without RBD but with Rfam ID (light green), non-RBPs with RBD and Rfam ID (dark green), RBPs without RBD nor Rfam ID (light orange), RBPs with RBD but without Rfam ID (dark orange), RBPs without RBD but with Rfam ID (light purple) and RBPs with RBD and Rfam ID (dark purple). *, ** and *** correspond to *P*-values < 0.05, 0.01 and 0.001, respectively, ns corresponds to non-significant, as resulting from a Wilcoxon rank sum test.

### Retrieval of the MobiDB-lite IDR content fraction

Among the various algorithms that were available for the prediction of IDRs (39), we selected the MobiDB-lite prediction as it is more conservative, through the combination of 10 different prediction tools (34,39). To retrieve the IDR content fraction of the proteins, we downloaded the results of the MobiDB-lite algorithm from the MobiDB database (https://mobidb.bio.unipd.it, Version 5.0.0) (34,40) for each of the 13 RBP2GO species. The Supplementary Table S4 was accordingly updated with this information (IDR content fraction).

The results of these analyses were used to produce Figure 5A and Supplementary Figure S6 and S7.

**Figure 5.**
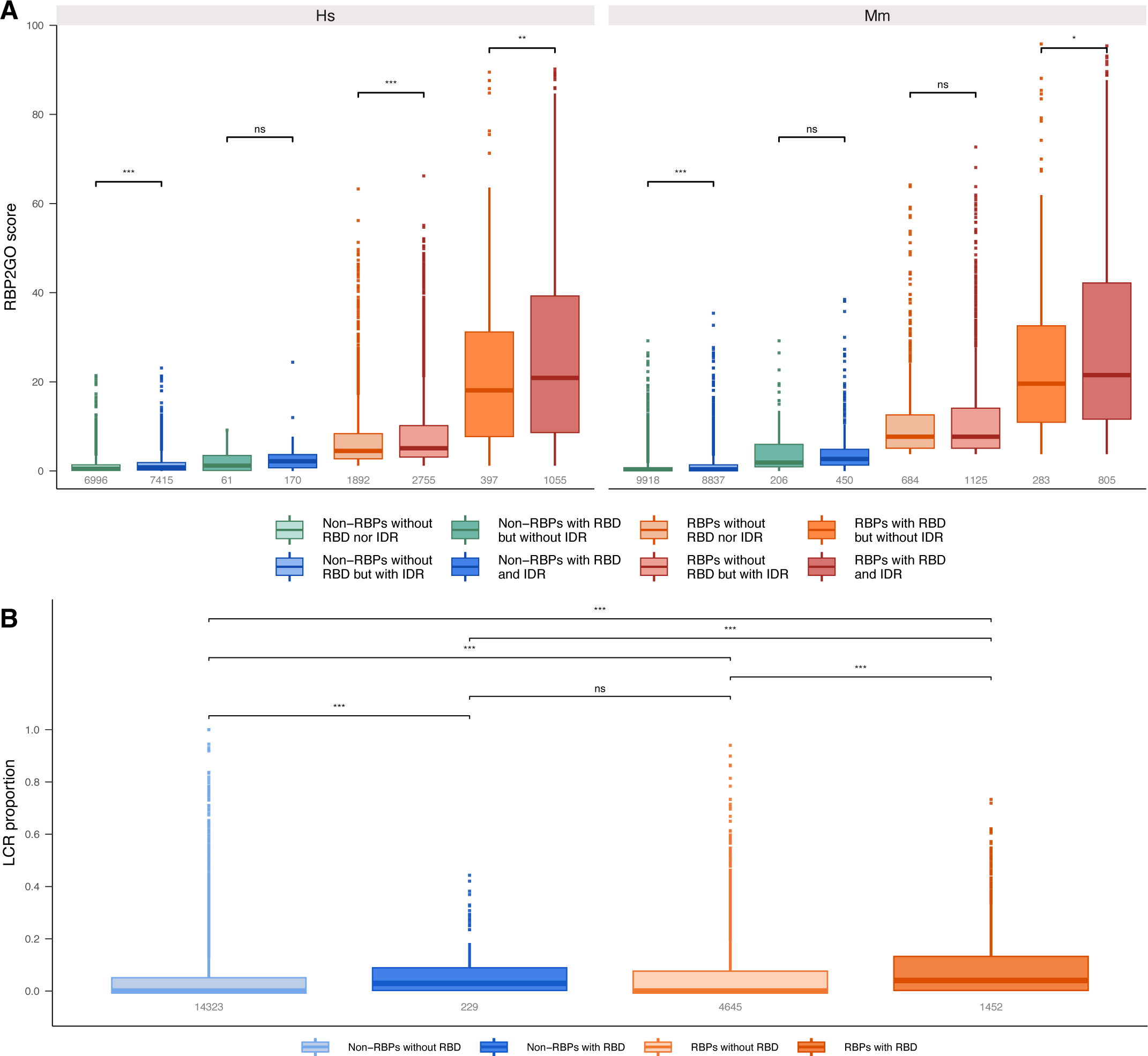
Analysis of the disordered fraction in the human and murine proteins of the RBP2GO database. (**A**) Boxplots representing the distribution of RBP2GO score in eight groups of proteins: non-RBPs without RBD nor IDR (light green), non-RBPs without RBD but with IDR (light blue), non-RBPs with RBD but without IDR (dark green), non-RBPs with RBD and IDR (dark blue), RBPs without RBD nor IDR (light orange), RBPs without RBD but with IDR (light red), RBPs with RBD but without IDR (dark orange) and RBPs with RBD and IDR (dark red). (**B**) Boxplots representing the distribution of LCRs in non-RBPs without RBD (light blue), non-RBPs with RBD (dark blue), RBPs without RBD (light orange) and RBPs with RBD (dark orange). *, ** and *** correspond to *P*-values < 0.05, 0.01 and 0.001, respectively, ns corresponds to non-significant, as resulting from a Wilcoxon rank sum test.

### Linear regressions

For all linear regressions in this study, the regression lines were generated using the geom_smooth() function from the ggplot2 package (41), with the “lm” method. The correlation coefficients in the figures were computed using the R base function “cor()” or the “stat_cor()” function from the ggpubr package (42).

### Retrieval of the LCR frequency and amino acid compositional bias in the human proteins

LCRs and sequences of the proteins were obtained directly from the UniProt database using the UniProt.ws package (36): UniProt release 2023_05 from November 8^th^, 2023. These were used to compute the LCR proportion within the proteins of the four main groups (non-RBPs without RBD, non-RBPs with RBD, RBPs without RBD and RBPs with RBD) as depicted in Figure 5B. Amino acids were grouped in categories (-charge: negatively charged; + charge: positively charged; Aromatic, Uncharged non-polar and uncharged polar). The frequencies of specific motifs such as FGG, RGG and YGG were calculated based on the protein sequence and according to the four main groups of proteins.

The results of these analyses were used to produce the panels in Supplementary Figure S8.

### GO enrichment analysis for non-RBPs containing an RBD

For each species, a GO-enrichment analysis was performed in the non-RBPs containing one of the selected RBDs against the proteome with the PANTHER 17.0 classification system (http://pantherdb.org, release 2021_03) that was accessed via the “rbioapi” package in R (43). The results for the different species were then merged in a single table (Supplementary Table S7). The GO terms were filtered to keep the terms with an FDR <0.05. For each GO term, the number of species in which it was significantly enriched and its mean fold enrichment in these species were calculated. For the heatmaps (Figure 6), only the GO terms significantly enriched (FDR < 0.05) in more than four species out of 13, and with a mean fold enrichment > 5, were selected. The heatmaps were created with the geom_tile() function from the ggplot2 package (41), and the GO terms were sorted by decreasing mean fold enrichment (Figure 6).

**Figure 6.**
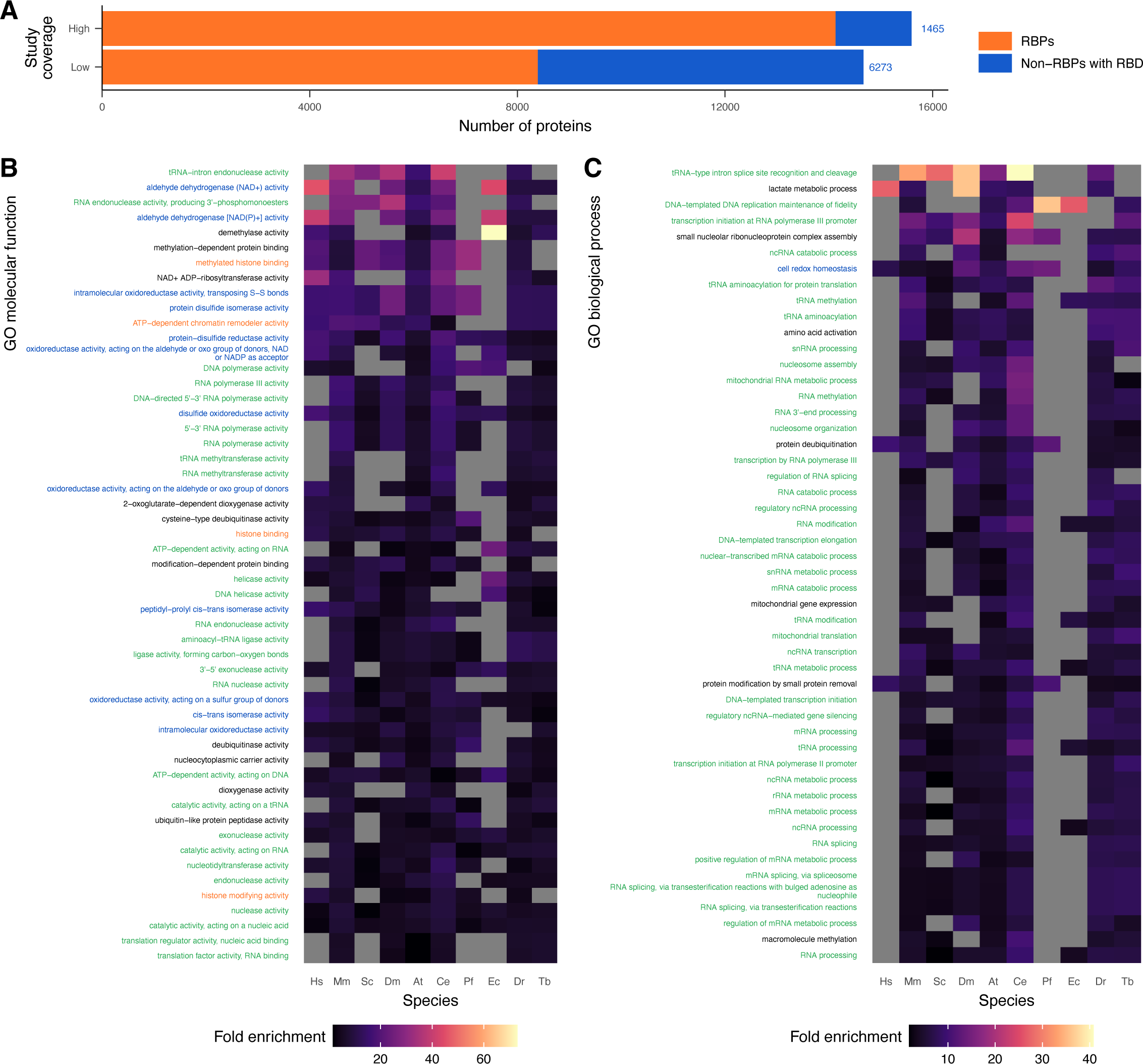
Discovery of candidates for RBP aspirants using the RBD content of the non-RBP proteins. (**A**) Number of RBPs and RBD-containing non-RBPs (numbers indicated in blue for this group) for species with a high (≥9, top bar) or a low (<9, bottom bar) number of RBP screen studies. (**B**) Heatmap showing the most enriched GO terms for molecular function in the RBD-containing non-RBPs as compared to the proteome of each species. The GO terms were classified by mean fold enrichment, from top to bottom. The terms are related among other molecular functions to metabolism (blue), to DNA and RNA (green), and to chromatin regulation (orange). (**C**) Same as in (**B**) for the most enriched GO terms for biological processes.

### Identification of new RBDs using the human RBPs lacking canonical RBDs

In this part of the analysis, we only included the human proteins as it is the species covered by the highest number of datasets. Based on their RBP2GO score, we selected the top 20% of the RBPs that did not contain any of the selected RBDs (935 “high score” proteins of interest, Supplementary Table S4) that we compared to three reference datasets: 1) the whole proteome (20741 proteins), 2) the non-RBPs without RBD (14411 proteins) and 3) the bottom 20% RBPs lacking RBDs based on their RBP2GO score (989 “low score” proteins, Supplementary Table S4). We extracted the InterPro IDs present in the “high score” dataset and filtered out the IDs that were present in the initial RBD candidate list. Next, we selected for the IDs of the type “Domain” or “Repeat”, which resulted in a list of 637 IDs (Supplementary Table S8). For each InterPro ID (domain/repeat), we calculated the proportion of proteins containing the domain in the dataset of interest as well as in the three reference datasets. Next, for each domain, we computed its fold enrichment in the dataset of interest as compared to each of the reference datasets. A *P*-value was also generated as a result of a Fisher’s exact test. The *P*-values were further adjusted for multiple testing by the FDR method (44). To identify new RBDs, we applied the following criteria: we selected the domains that were 1) significantly enriched against at least one reference dataset (>2-fold enrichment and adjusted *P*-value <0.05), 2) present in at least 0.5% of the proteins of the “high score” dataset (*i.e.*, in at least five proteins) and 3) presenting an RBP vs. non-RBP hits ratio greater 1. Altogether, 15 domains satisfied these three criteria (Supplementary Table S8).

The results of these analyses were used to produce the panels A to E in Figure 7.

**Figure 7.**
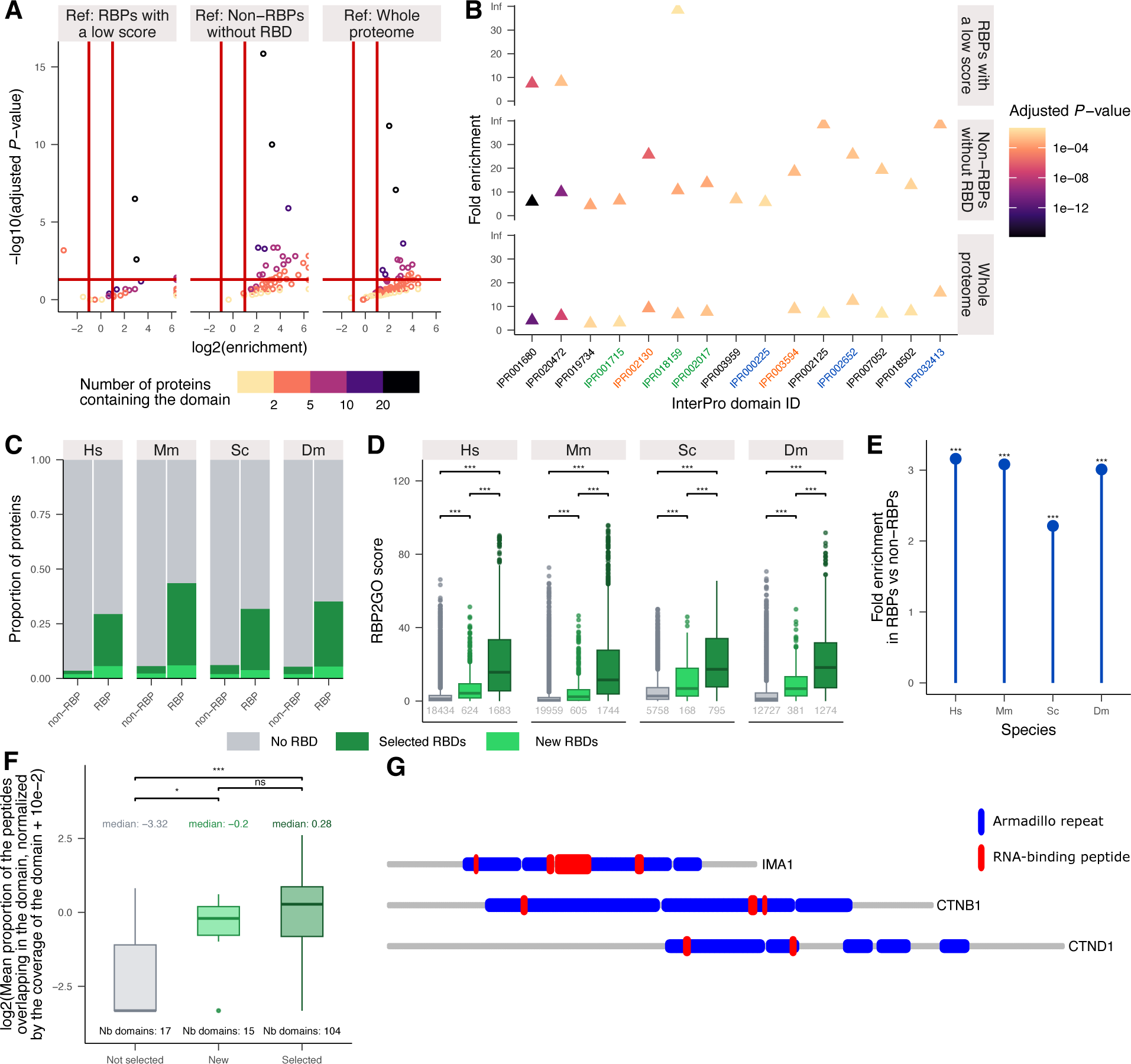
Discovery of new RBD candidates in RBPs lacking RBDs. (**A**) Volcano plots showing the enrichment and its adjusted *P*-values for each of the InterPro domains present in the RBPs without RBD within the top 20% of RBP2GO scores, with an RBP/non-RBP ratio > 1, as compared to three different control groups (RBPs without RBD within the bottom 20% RBP2GO scores; non-RBPs without RBD; whole proteome). The *P*-values were calculated using a Fisher’s exact test and were adjusted with the FDR method. (**B**) Enrichment and *P*-values for the 15 InterPro IDs selected as candidate RBDs, see also Supplementary Table S8. Some InterPro IDs are related to chaperones (orange), cytoskeletal-protein binding (green), or importins (blue). (**C**) Proportion of proteins without RBD (grey), with selected RBDs (dark green) or new RBD candidates (light green) in non-RBPs or RBPs. (**D**) Boxplots showing the distribution of the RBP2GO score for the proteins without RBD (grey), containing a new RBD candidate (light green), or a selected RBD (dark green). *** corresponds to *P*-values < 0.001 as resulting from a Wilcoxon rank sum test. The numbers given in grey represent the size of the protein groups. (**E**) Fold enrichment of RBPs over non-RBPs across all proteins containing one of the new RBD candidates. The *P*-values were calculated using a Fisher’s exact test. (**F**) Boxplots showing the distribution of the log2 of the normalized mean proportion of the RNA-binding peptides overlapping the domain + 1×10^-2^. The domains were categorized as not selected RBDs (grey, see Figure 1), new RBD candidates (light green) and selected RBDs (dark green, see Figure 1). Only domain present in at least 5 proteins were taken into account (see Material and Methods section for more details). The number above each box represents the median values and the number under each box represents the size of the group. The *P*-values were calculated using a Wilcoxon rank sum test. (**G**) Schematic representations of three proteins (grey) with Armadillo repeats (blue) and the overlapping RNA-binding peptides (red).

### Validation of the newly discovered RBDs using RNA-binding peptides

Several studies featured in RBP2GO used approaches enabling the identification of the RNA-binding region of the detected RBPs (13,14,16,32,45–47). Using these datasets, we computed a new dataset combining the peptide information with the UniProt ID of the corresponding proteins and with the coordinates of the peptides in the proteins (Supplementary Table S9). Hereby, we only considered the human protein datasets. For each of the 624 proteins containing only the new RBDs (without containing any of the previously selected RBDs), we computed the proportion of peptides overlapping at least 50% with the new RBD relative to the total number of peptides found in the protein. This number was normalized to the fraction of the protein sequence that was covered by the domain. We then summed this number for all the domains corresponding to one InterPro ID over all proteins containing at least one peptide, and normalized by the total number of occurrences of this domain in the proteins containing at least one peptide. The same procedure was repeated for the selected RBDs and the non-selected domains (Figure 1A) using groups of proteins containing only selected RBDs or only non-selected domains respectively. To maintain the same selection criteria as for the new RBDs, we only took into account the selected and non-selected RBDs that were present in at least five RBPs (Supplementary Table S9). The results of these three analyses were used to produce Figure 7F-G.

### Computation of the RBP2GO composite score

To reflect the importance of the presence of RBDs or the relation to an Rfam ID for an RBP, we created a new “RBP2GO composite score” that integrates the knowledge on RBDs and Rfam IDs in addition to the information already provided by the RBP2GO score. This new score comprises three components. Component 1 is based on the listing count of the proteins (i.e., in how many datasets the protein is detected as RBP). Component 2 is based on the average listing count for the top 10 interactors of the protein, according to the STRING database (STRING-DB.org) (48). Component 3 provides a score pertaining to the presence of RBDs or Rfam IDs that are associated to the protein of interest.

To define the component 3, a quality factor was attributed to each RBD / Rfam ID. To this aim, we took into consideration the ratio of hits in RBPs versus hits in non-RBPs in human (Hs) as a surrogate measure for the likelihood of the RBD or Rfam ID being truly associated with RBPs. If the ID was not found in Hs, we considered the ratio in mouse (Mm); if it was not found in Mm, we considered the one in yeast (Sc), and so on following the order of the species by decreasing number of studies (Supplementary Figure S1). For each ID, this ratio was used to attribute a semi-quantitative balancing quality factor: ratio <2 = quality factor of 1; ratio ≥2 = quality factor of 2; ratio ≥4 = quality factor of 3; ratio ≥8 = quality factor of 4 and ratio ≥16 = quality factor of 5. For infinite ratios (ID only present in RBPs), the number of RBPs with this ID was in addition taken into account: for a number of RBPs ≤2 = quality factor of 2; number of RBPs from 2 to 4 = quality factor of 3; number of RBPs from 4 to 8 = quality factor of 4 and number of RBPs >8 = quality factor of 5. The attribution rules are summarized in the Supplementary Table S10. Finally, we summed the quality factors for all RBDs and RNA-related family IDs present for one protein, and limited this number to 25.

In order to weight properly the contribution of each score, we applied a Random Forest algorithm in R, using three independent datasets of RBPs and non-RBPs. Dataset 1: based on the proteins of the RBP2GO database (22); dataset 2: based on the HydRA classification (49); dataset 3: based on RBPbase superset (3). As a result of the Random Forest analysis and the calculated importance of the components, while the RBP2GO composite score overall ranges from 0 to 100, component 1 can amount to a maximum of 50, while component 2 and 3 can equally amount to a maximum of 25 (Figure 8 and Supplementary Table S11).

**Figure 8.**
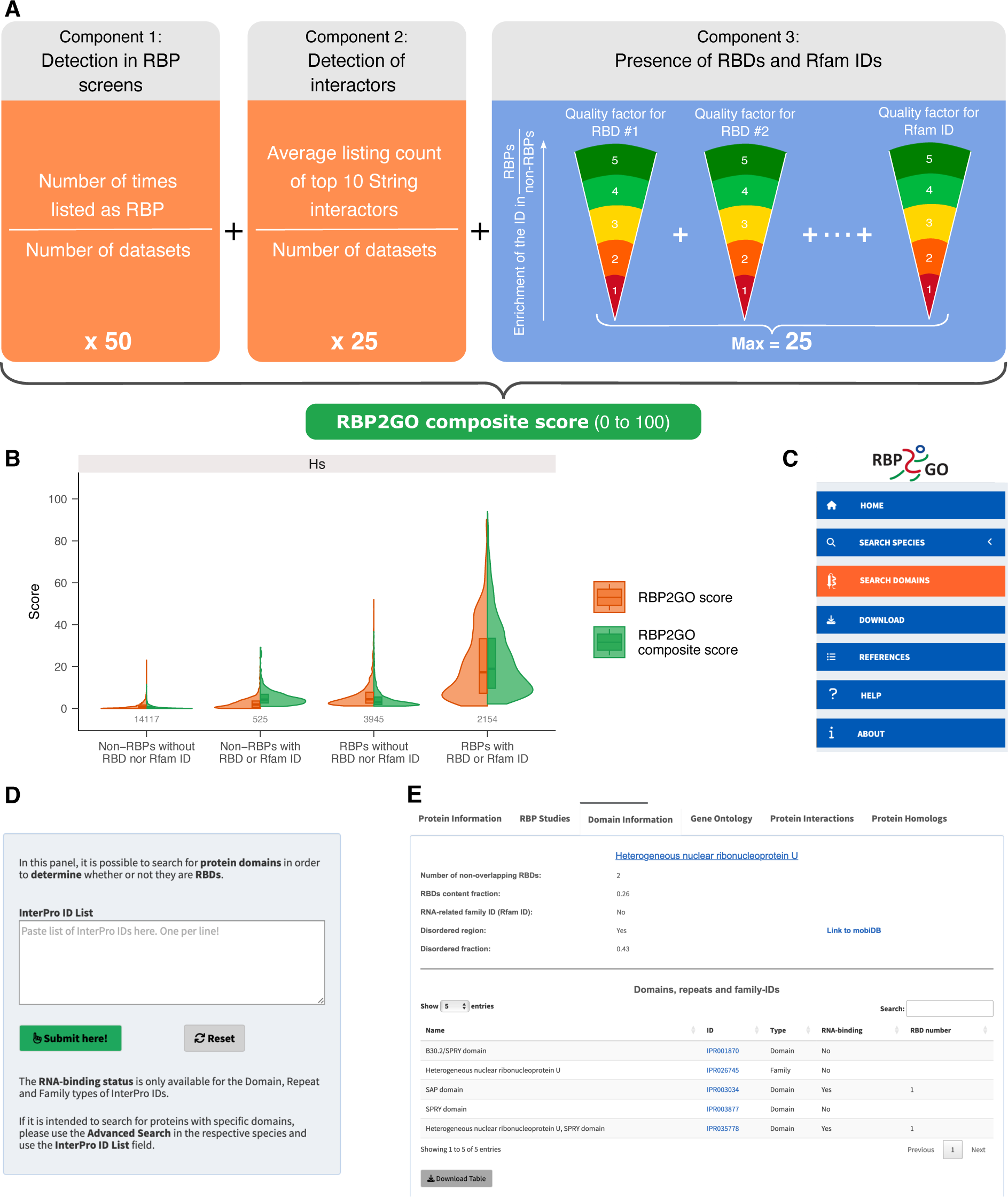
Development of the RBP2GO composite score and update of the RBP2GO database. (**A**) Schematic representation of the compilation of the RBP2GO composite score. The quality factors as well as their attribution rules are explained in the Supplementary Table S10. The maximum values are 50 for component 1 and 25 for each component 2 and 3. The RBP2GO composite score takes values between 0 and 100. (**B**) Violin plots representing the distribution of the RBP2GO score (orange) and the RBP2GO composite score (green) for non-RBPs without RBD nor Rfam ID, for non-RBPs with RBD and/or Rfam ID, for RBPs without RBD nor Rfam ID and for RBP with RBD and/or Rfam ID. (**C**) Screenshot of the sidebar of the advanced RBP2GO database (https://RBP2GO-2-Beta.dkfz.de) showing the added menu for domains and families in orange. (**D**) Screenshot of the additional search function dedicated to the protein domains (found under the SEARCH DOMAINS of the sidebar as shown in **C**). (**E**) Screenshot of the new “Domain information” tab added to the protein search section of each species in the RBP2GO database. This new tab displays all information pertaining to the RBDs and Rfam IDs of the selected protein, as well as information on its disorder content. It includes links to the InterPro (33) and MobiDB (40) databases. For each InterPro ID, a table indicates the type of the ID as “Domain”, “Repeat” or “Family” and if it is linked to “RNA-binding”.

The Pearson correlation coefficients between the three components of the composite score and the RBP2GO score were computed using the cor() function of the stats package in R. The results are displayed in Supplementary Figure S9.

The RBP2GO composite score was analyzed in the four groups of proteins, *i.e.*, non-RBPs without RBD, non-RBPs with RBD, RBPs without RBD and RBPs with RBD to produce the panel B in Figure 8. Supplementary Table S4 was updated accordingly.

As an application of the RBP2GO composite score, the various proteome-wide strategies were grouped according to their underlying methods, and the identified RBPs were analyzed, according to the three components and the RBP2GO composite score itself (Supplementary Figure S10 and Supplementary Table S12).

### Selection of high-confidence RBPs in *Homo sapiens* (Hs) and comparison with HydRA and RBPbase RBPs

We took advantage of the RBP2GO composite score to compute a list of selected high-confidence human RBPs. This list encompassed all human proteins, whether they were detected as RBPs in proteome-wide screens or not, that had a composite score superior or equal to 10, with at least two out of the three components of the score larger than zero. This resulted in a list of 2018 proteins, among which 1977 were already detected as RBPs in screens of the RBP2GO database (Supplementary Table S13, Supplementary Figure S11). We compared the high-confidence RBP assignment to other approaches, such as HydRA and RBPbase. The distributions of the RBP2GO composite scores and HydRA scores were calculated for proteins with both scores available (Supplementary Figure S12).

### Update of the RBP2GO database

The whole dataset compiled in this analysis is available as Supplementary Table S14. In order to make the data more accessible and searchable, this new dataset was included in a user-friendly and intuitive manner into the RBP2GO database. The “RBD_Status” column as well as the number of RBDs and the RBD content fractions were updated to take into account the identified RBDs (Supplementary Table S8) in addition to the selected RBDs and Rfam IDs. The new data, including the lists of RBDs and Rfam IDs can be directly downloaded from the download section of the database (https://RBP2GO-2-Beta.dkfz.de). It is now also possible via the advanced search option from each species to search for proteins containing a specific InterPro ID, or containing an RBD and/or an Rfam ID. A new section with information about the selection procedures applied to the RBDs and Rfam IDs was also created under the sidebar menu “search domains”, which include a specific search module for InterPro IDs.

For easy access to the analyze and the datasets, the whole script is available from the download section of the RBP2GO database from the sidebar menu.

## RESULTS

### Proteome-wide RBP screens allow refining the pool of RNA-binding domains

Prior to the increase in the number of proteome-wide methods dedicated to the identification of RBPs and the resulting knowledge gained regarding RNA-protein interactions, RBPs were thought to interact with RNA essentially via well-defined globular RBDs. Some proteins harbor a single RBD while others contain several RBDs, combined in a modular manner, which illustrates the great variability of RBD-RNA interactions (24,26).

Based on the datasets compiled in our recently published RBP2GO database (22), RBP candidates were defined as proteins that were at least detected in one RNA-interactome dataset (Listing Count >= 1 in the RBP2GO database). RBP candidates are thus not necessarily validated experimentally. For more simplicity, RBP candidates are named RBPs in the following. Non-RBPs were defined as the proteins that had never been identified in the proteome-wide experimental studies (Listing Count = 0). For each protein, we also defined an RBP2GO score, that reflects two aspects: first, we reasoned that proteins that are independently listed in multiple datasets must have higher probabilities of being true RBPs. Second, based on the finding of the SONAR study (25) and our previous analysis of the protein– protein interactions within the CORUM database (19), proteins interacting with multiple RBPs are frequently RBPs themselves. Accordingly, we computed and provided for each protein two separate indicators of its RBP propensity: the count of the protein itself being listed as RBP (Listing Count) and the average listing count of the up to ten interaction partners with the highest STRING interaction database scores (AVG10 Int Listing Count). These two components were combined with an equal weight in the RBP2GO score, which was then normalized to the number of datasets in the respective species. As a result, the RBP2GO score can range from 0 to 100.

Here, we used this knowledge on RBPs to refine the pool of RBDs. We combined and filtered RBD lists from previous works (8,13,29) together with data from the InterPro database (33) to compile a list of in total 2712 InterPro IDs corresponding to classical and non-classical RBDs (8) as well as domains with descriptions and GO terms related to RNA binding (Supplementary Table S1). By filtering for the “domain” and “repeat” types of InterPro IDs, we computed a starting list of 1289 RBD candidates (Figure 1A). Next, we applied a three-step selection procedure (as described in the Material and Methods section) that was based on the ratio between the presence of the domains in RBPs as compared to non-RBPs in the 13 species included in the RBP2GO database in the order of the number of available datasets for the respective species (Supplementary Figure S1): *Homo sapiens* (Hs), *Mus musculus* (Mm), *Saccharomyces cerevisiae* (Sc), *Drosophila melanogaster* (Dm), *Arabidopsis thaliana* (At), *Caenorhabditis elegans* (Ce), *Plasmodium falciparum* (Pf), *Escherichia coli* (Ec), *Danio rerio* (Dr), *Trypanosoma brucei* (Tb), *Salmonella Typhimurium* (ST), *Leishmania mexicana* (Lm) and *Leishmania donovani* (Ld) (22). This analysis resulted in a set of 977 selected RBDs, further referred to as RBDs in the following (Supplementary Table S3).

We evaluated this selection of RBD candidates using the RBP2GO score of the proteins containing the selected RBDs in species with a higher number of proteome-wide RBP datasets such as *Homo sapiens*, *Mus musculus*, *Saccharomyces cerevisiae*, *Drosophila melanogaster* (Supplementary Figure 1), hereafter also referred to as the “well-studied species”. We confirmed that these proteins had a significantly higher RBP2GO score than the proteins with unselected or no RBDs, which corroborated our selection procedure (Figure 1B).

### The proportion of RBD-containing proteins in proteomes differs across species

Next, we analyzed the proportion of the selected RBDs in the proteins of the RBP2GO database that were classified as RBP (Listing Count >= 1) or non-RBP (Listing Count = 0). Accordingly, the protein can be grouped into RBPs with RBD (RBPs containing at least one selected RBD), RBPs without RBD, non-RBPs with RBD and non-RBPs without RBD. While some well-studied species such as human and mouse (Supplementary Figure S1) harbored more than 65% of the RBDs only in the RBPs (Figure 2A and Supplementary Table S15), this seemed to be inverted in less studied species. In zebrafish (*Danio rerio*), 72% of the RBDs were found only in non-RBPs (Figure 2A and Supplementary Table S15).

This disparity between well-studied species and less-studied species was also reflected in the proportion of RBPs and RBD-containing proteins in the proteomes. The *Homo sapiens* proteome contained 3.5-fold less RBD-containing proteins than RBPs, while *Danio rerio* and *Trypanosoma brucei* inversely depicted >4.5-fold more RBD-containing proteins than RBPs (Supplementary Table S16). However, in the group of less-studied species, *Escherichia coli*, which had 5-fold more RBPs than RBD-containing proteins was an exception (Figure 2B).

Interestingly, in the well-studied species, while the proportion of RBPs in the proteome was higher than for most of the less-studied species, only a small proportion of the RBPs contained an RBD (<10% of the proteome, Figure 2C, dark orange group of proteins and Supplementary Table S16). Most RBPs did not contain an RBD, which strongly indicates that RBPs employ alternative ways in addition to already known RBDs to interact with RNA molecules.

The four most abundant RBDs in the RBPs of our dataset appeared all to be canonical RBDs (26,28), namely, the RNA recognition motif domain (RRM, IPR000504, found in 1700 proteins), Helicase superfamily 1/2 ATP-binding domain (IPR014001, found in 1079 proteins), Helicase C-terminal (IPR001650, found in 1060 proteins) and DEAD/DEAH box helicase domain (IPR011545, found in 701 proteins, Supplementary Table S4). Here again, there was a clear discrepancy between the well-studied species and less-studied species with regard to the experimental identification of proteins containing such domains as RBPs. In the well-studied species, more than 50% of the proteins harboring at least one of these four canonical RBDs were classified as RBPs (Figure 2D and Supplementary Table S4). For many less-studied species, the percentage of RBPs dropped down to less than 15% of the proteins. Thus, the proteomes of the less-studied species may contain a number of yet undiscovered RBPs, including proteins containing selected RBDs (Supplementary Table S3).

In humans, 231 RBD-containing proteins had not been detected yet in the published RBP screens (Supplementary Table S16). We hypothesized that the expression levels of the proteins might impact their potential of being detected in the mass spectrometry-based studies. Therefore, we analyzed the protein expression levels (37) of the four different groups: non-RBPs without RBD, non-RBPs with RBD, RBPs without RBD and RBPs with RBD. Both non-RBP groups had significantly lower expression levels (Figure 2E), which might potentially explain why the non-RBPs with RBD had not been detected as RBPs yet. Similarly, RBP2GO scores, which reflect the probability for a protein to be an RBP (22), positively correlated with the protein expression levels (Supplementary Figure S2).

In summary, a substantial number of non-RBPs with RBD, further referred as “RBP aspirants”, had not been detected in proteome-wide screens so far - especially in the less-studied species. Furthermore, the high proportion of RBPs without RBD, especially in the well-studied species, raises the question of how these proteins interact with RNA.

### Proteins with RBD appear as stronger RBPs

Taking advantage of the RBP2GO score (22), we evaluated its distribution in the different species and found that RBD-containing proteins had a significantly higher RBP2GO score in all species as compared to proteins without RBD (Figure 3A and Supplementary Figure S3A). This was confirmed when the non-RBPs and RBPs were split into two groups, depending on their RBD content. In each group respectively, the RBD-containing proteins had significantly higher RBP2GO scores, with the exception of *Danio rerio* and *Trypanosoma brucei* (Figure 3B and Supplementary Figure S3B). This was, however, linked to the low number of proteins detected as RBPs so far in these two species (Figure 2B and Supplementary Table S16).

To obtain a more detailed view of the interplay between the presence of RBDs in proteins and the RBP2GO score, we evaluated the proportion of RBD-containing proteins for each interval of the RBP2GO score. We found a strong positive correlation between the proportion of RBD-containing proteins and the RBP2GO score up to a score of 50 (Supplementary Figure S3C). Above 50, for the four well-studied species and *Caenorhabditis elegans*, >80% of the proteins contained an RBD (Supplementary Figure S3D and Supplementary Table S17). This result supports the assumption that RBDs can be used to assess the RNA-binding potential of an RBP, but also to identify new RBP candidates in the non-RBPs.

Exploiting detailed information from the InterPro database (33), we compiled the number of RBDs per protein, as well as the fraction of the protein sequence they covered. We analyzed this data in RBPs versus non-RBPs and observed that RBPs had a higher RBD content fraction than non-RBPs, a difference that was significant for all species except *Plasmodium falciparum*, *Escherichia coli* and *Trypanosoma brucei* (Figure 3C and Supplementary Figure S3E). Moreover, in human proteins, the RBP2GO score increased with the number of RBDs per protein, whereas this was not visible for the non-RBPs (Figure 3D).

Taken together, these results highlight the strong positive correlation between the RBP2GO score and the number of RBDs in RBPs. This underlines the potential of RBDs for the evaluation of the RNA-binding ability of proteins, including non-RBPs with RBD, that might not be properly detected if lowly expressed.

### The RBP-enriched Rfam IDs are also strong indicators of RNA-binding capacity

In addition to the information relative to the protein domains and repeats, the InterPro database provides annotation for protein families. In the initial lists of RNA binding-related InterPro IDs, 1028 IDs corresponded to “family” IDs (Supplementary Table S1). We subjected these IDs to the selection procedure, which we previously applied to the “domain” and “repeat” IDs (Figure 1A and Supplementary Figure S4A). We obtained a list of 672 selected RNA-related “family” IDs, referred to as Rfam IDs (Supplementary Tables S6 and Supplementary Figure S4A). The distribution of the Rfam IDs across RBPs and non-RBPs in the 13 RBP2GO species exhibited a similar pattern as for the RBDs (Figures 2A and 4A). The well-studied species displayed a large proportion of the Rfam IDs in RBPs only. 98.2% of the Rfam IDs in *Homo sapiens* and more than 65% in *Mus musculus* were found in RBPs. In less-studied species such as *Trypanosoma brucei*, less than 7% of the Rfam IDs were related to RBPs only (Figure 4A and Supplementary Table S18). These observations confirmed the importance of a good study coverage for proper classification of proteins into RBPs or non-RBPs. Strikingly, 70% of the human RBPs were not linked to RNA-related InterPro ID (Supplementary Figure S4B).

The proteins annotated with an Rfam ID had a significantly higher RBP2GO score than the ones without Rfam ID (Figure 4B and Supplementary Figure S5A). This was also verified when splitting the proteins into non-RBPs and RBPs (Supplementary Figure S5B-C), except for the human non-RBPs, for which the difference was not significant mainly due to the small number of non-RBPs with Rfam ID.

There was also a strong positive correlation between the RBP2GO score and the proportion of proteins associated with Rfam ID (Supplementary Figure S5D). For the well-studied species and within the proteins with RBP2GO scores >50, proteins with RBD were more abundant than proteins with Rfam ID (Supplementary Figure S3A and S5D). However, proteins with Rfam ID annotations seemed to have higher median RBP2GO scores than proteins with RBD only (Figure 4C and Supplementary Figure S5F). In addition, in five out of nine species, RBPs annotated with both, RBDs and Rfam IDs, had a significantly higher RBP2GO score than the other RBP groups, indicating a cumulative effect of the dual annotation (Figure 4C and Supplementary Figure S5F).

Altogether, these results demonstrate that the selected Rfam IDs, similar to the RBDs, represent important protein features that can improve the evaluation of the RNA-binding potential of RBPs. While annotations with either RBDs or Rfam IDs seemed equivalent, having both annotations was linked to a higher RBP2GO score in several species, hence a greater likelihood for the protein to harbor physiologically relevant RNA-binding abilities.

### Disordered regions are enriched in proteins with RBD

While RBDs and Rfam IDs were both strong indicators of the RNA-binding potential of proteins, our analysis revealed a large group of RBPs (*e.g.,* 70% in human) that were lacking such features (Supplementary Figure S4B and Supplementary Table S16). In the past decades, protein regions among those regions referred to as IDRs were experimentally detected as directly interacting with RNA (13,32). IDRs are defined according to their structural properties. Unlike globular domains, IDRs do not fold into a specific three-dimensional structure, but rather remain lose and confer flexibility to proteins (50). Their structure can stabilize upon binding to interaction partners (51). Some IDRs are implicated in RNA binding for example via specific repeats such as the arginine and serine (RS) repeats in the NF-kappa-B-activating protein (NKAP), which plays an important role in splicing (30,31). Also, IDRs that contain repeats of arginine and glycine, referred to as RGG-box, can also bind directly to RNA. Such IDR repeats are for example found in the fragile X mental retardation protein (FMRP/FMR1) (30,52). This motivated us to analyze the IDR content of the proteins.

To do so, we retrieved the data computed by the MobiDB-lite algorithm (34,40). The analysis of the IDR content of the proteins revealed that RBD-containing proteins had a significantly higher IDR fraction than proteins lacking RBDs, regardless of their status as RBP or non-RBP, in most species but especially in the well-studied species (Supplementary Figure S6). This is consistent with previous findings reporting the enrichment of IDRs in RBPs (13,53). In most species, the RBPs without RBD did not exhibit a higher disordered fraction than the RBD-containing non-RBPs, indicating that their RNA-binding ability was probably independent of the IDR content (Supplementary Figure S6).

The combined analysis of the RBP2GO score of the RBPs with respect to the RBD and the IDR content revealed that the presence of IDRs in addition to RBDs only had a minor positive contribution to the increase of the RBP2GO score (Figure 5A and Supplementary Figure S7A). The minor contribution of the IDR to the RBP2GO score of the proteins was further validated by the weak linear correlation between the disordered fraction and the RBP2GO score for both the RBPs and the non-RBPs (Supplementary Figure S7B).

According to our results, the IDRs appeared enriched in proteins that already contained RBDs (both RBPs and non-RBPs). Therefore, the presence of IDRs in proteins seems largely redundant with the presence of RBDs and does not represent a strong independent indicator of the RNA-binding potential of proteins that would otherwise explain, for example, how RBPs lacking RBDs could bind RNA.

IDRs frequently contain repeats, i.e., the underlying sequences depict amino acid content bias. Such sequences are also referred to as low complexity sequences or low complexity regions (LCRs). Since specific LCRs can bind to RNA (31,52), we also investigated the LCR content of the human RBPs and non-RBPs. We found that charged amino acids were slightly enriched in RBPs (Supplementary Figure S8A). More specifically, we observed that RGG and YGG repeats were enriched in proteins with RBD, suggesting a co-occurrence of LCRs with RBD (Supplementary Figure S8B), similar to the co-occurrence of IDRs with RBD. In support of this finding, human proteins with RBD (both non-RBPs and RBPs) had a significantly higher proportion of LCRs than the proteins lacking RBDs (Figure 5B).

### New RBP candidates can be identified from non-RBPs with RBD

When splitting the 13 studied species into well-studied and less-studied species according to the number of available datasets (Supplementary Figure S1), the four well-studied species summed up together 1465 non-RBPs with RBD out of 5496 proteins with RBD (26.6%). For the nine less-studied species, there were 6273 non-RBPs with RBD out of 9308 proteins with RBD (67.3%) (Figure 6A and Supplementary Table S16). Adding the non-RBPs with RBD to the collection of RBPs would increase the set of RBPs by 75% in the less-covered species as compared to only 10% in the well-studied species.

In order to evaluate the potential of the non-RBPs with RBD as new RBP aspirants, we performed a Gene Ontology (GO) (54) enrichment analysis on the non-RBPs with RBD in the eight species for which data was available from the PANTHER resource (55,56). We report the terms that were enriched in at least half of the species, sorted by the mean fold enrichment (Figure 6B-C). In the enriched GO terms for molecular function and biological processes, many terms were related to nucleic acid binding and metabolism such as “RNA polymerase activity” (as depicted in green in Figure 6B-C). In agreement with these findings, several high-mobility group (HMG) chromosomal proteins were already listed as RBPs in the RBP2GO database (22) and experimentally validated, i.e., HMGN1 (19). Also, DNA-binding domains such as the high-mobility group (HMG)-box had already been associated to RNA-binding (13,57). Inversely, heterogeneous ribonucleoproteins and double-stranded RNA-binding proteins could also directly interact with DNA (58,59), illustrating that many proteins binding to DNA can also bind to RNA and *vice versa* (12,60). GO terms related to histones and chromatin were enriched, as well, but were less represented (orange, Figure 6B-C). RBPs can interact with chromatin and act in gene regulation, as for example the RNA-binding protein 25 (RBM25) which enhanced the transcription-dependent activities of the YY1 transcription factor (61–63). Metabolic activities also appeared to be enriched in the non-RBPs with RBD (blue, Figure 6B-C). This is also in agreement with the identification of several metabolic enzymes in RBPome studies and their further validation such as the iron regulatory protein 1 (IRP1), the glyceraldehyde-3-phosphate dehydrogenase (GAPDH) and the glycolytic enzyme Enolase 1 (ENO1)(64–67). Altogether, our GO analysis of the non-RBPs with RBD strongly indicated that this group was enriched in molecular functions and biological processes reasonably linked to RNA such that this group may contain further so far experimentally unrecognized RBPs, referred to as RBP aspirants.

To further validate these findings, we found evidence in the literature for the RNA-binding properties of seven RBD-containing non-RBPs, which contained different types of RBDs (Supplementary Table S19). Individual studies experimentally confirmed the binding of these proteins to RNA (68–73), in support of our assumption that RBP aspirants (non-RBPs with RBD) are indeed RBPs. Furthermore, six out of these seven proteins exhibited IDRs, in agreement with the frequent association of IDRs and RBDs (Supplementary Table S16) as suggested by the enrichment of IDRs in RBPs with canonical RBDs (74) and confirmed in our analysis above (Figure 5A).

We showed here that a substantial number of non-RBPs with RBD, herein called RBP aspirants, are probably RBPs that had not been detected in proteome-wide screens so far. Using our selected list of RBDs as indicator of their RBP potential allowed us to identify 7738 new RBP aspirants, with more than 80% of them in less-studied species.

### New RBDs can be predicted in RBPs without known RBDs and high RBP2GO score

Since a large proportion of RBPs did not contain any of the selected RBDs, we hypothesized that they may contain so far undetected RBDs. To test this hypothesis, we performed an enrichment analysis of the “domain” and “repeat” InterPro IDs present in the top 20% of the human RBPs without RBD based on their RBP2GO score. This group of 935 proteins was compared to three different reference sets: the bottom 20% of the human RBPs based on their RBP2GO score, the non-RBPs without RBD and the human proteome (Figure 7A and Supplementary Table S4). 15 domains were significantly enriched as compared to at least one of the reference datasets. Within these domains, we found domains related to chaperones and importins, along with domains related to the binding of cytoskeletal proteins (Figure 7B and Supplementary Table S8). In particular, the WD40 repeat (IPR001680) does not exhibit a role in a specific cellular process, but is found in RBPs such as the WD-repeat containing protein 43 (WDR43) (75). The histidine kinase/HSP90-like ATPase domain (IPR003594) is present in the HSP90A protein which binds to RNA and is implicated in the loading of small RNAs into the RISC complexes (76). These findings could corroborate the RNA-binding ability of the 15 identified RBD candidates. In addition, a literature search based on these 15 RBD candidates revealed that the WD40 repeat (IPR001680) indeed is directly binding to RNA within the protein Gem-associated protein 5 or gemin5 (77), another evidence in support of the utility of our pipeline to identify RBDs among domains that were not yet annotated as such in the InterPro database. For the other 14 domains, we did not find RNA-binding evidence in the literature.

To validate the domains as RBDs, we quantified the proportion of proteins containing these domains. In the well-studied species, only a small proportion of RBPs harbored new RBD candidates as compared to RBPs with RBD (Figure 7C). In those species, the proteins with new RBD candidates had a significantly higher RBP2GO score than RBPs without RBD. However, their RBP2GO score was still significantly lower than that of the proteins with known RBDs (Figure 7D). Also, proteins containing only new RBD candidates were significantly enriched in RBPs as compared to non-RBPs (Figure 7E). These results supported the hypothesis that the new RBD candidates can be true RBDs.

To further validate the newly identified RBD candidates, we took advantage of RNA-binding peptide datasets available from human proteome-wide RBP studies (13,16,32,45). We calculated and normalized the proportion of these peptides overlapping the newly discovered RBDs, as compared to the overlap with RBD and InterPro domains that were not selected. The median proportion of overlapping peptides for the newly discovered RBDs appeared significantly higher than for the non-selected domains and not significantly different as compared to the selected RBDs (Figure 7F). For example, RNA-binding peptides were mapped to the Armadillo repeat (IPR000225), one of the newly identified RBDs (Figure 7G).

In summary, our pipeline identified 15 RBDs, of which 14 had not been previously linked to RNA. We validated those new RBDs using both the RBP2GO score of the related proteins as well as published RNA-binding peptide datasets. Consequently, we included them into the list of RBDs that can now be explored on the updated RBP2GO database (https://RBP2GO-2-Beta.dkfz.de).

### A new RBP2GO composite score helps defining a high-confidence human RBP list

In order to integrate the knowledge gained from the analysis of the RBDs and Rfam IDs, we computed a new RBP2GO composite score for each protein. This score combines three components that are based on experimental data directly related to the protein (component 1) and the top 10 interacting proteins according to the STRING database (component 2), as well as data related to RBDs and Rfam IDs (component 3, Figure 8A and Supplementary Table S10). Refer to the Material and Methods section for a detailed description. As expected, per definition, the first two components of the RBP2GO composite score strongly correlated with each other and with the RBP2GO score, but the correlation was decreased with the component 3 indicating that it added another layer of information (Supplementary Figure S9). The RBP2GO composite score allowed a better discrimination between proteins with and without RBD (Figure 8B, Supplementary Table S14), providing a solid support for the evaluation of RBPs. Moreover, while the previous RBP2GO score was only available for 9 out of 13 RBP2GO species due to a lack of study coverage (22), the component 3 of the RBP2GO composite score is now provided for all species.

The RBP2GO composite score can be used to learn about the different proteome-wide strategies used to identify RBPs with regards to the three components of the score. As depicted in Supplementary Figure S10, the various methods depict different spectrum of components and RBP2GO composite score. The distributions of the three components and the RBP2GO composite score present differences, from strategies with higher scores for all components, to strategies scoring better for components 1 and 2 than for component 3.

Finally, we took advantage of the RBP2GO composite score to define a high-confidence list of human RBPs that contains 2018 proteins with an RBP2GO composite score >10 and with at least two components >0 (Supplementary Table S13). As a validation of both the RBP2GO composite score and the high-confidence list, we found that human ribosomal proteins had a significantly higher RBP2GO composite score than all other proteins (Supplementary Figure S11) and were as expected all included in the high-confidence list of human RBPs (Supplementary Table S13). We also compared our high-confidence RBP assignment to the RBP assignment from HydRa (49) and the RBPbase superset (3). The intersection of the three datasets contains 1796 proteins, which represent 89% of the RBP2GO high-confidence human RBPs (Supplementary Figure S12A). We also found that the high-confidence human RBPs from our analysis had both a significantly higher RBP2GO composite score and HydRA score (Supplementary Figure S12B-C), confirming the quality of this list as a core catalog of human RBPs.

### An advanced version of the RBP2GO database includes detailed knowledge on RBDs and Rfam IDs

Our analysis demonstrated that RBDs and Rfam IDs were strong indicators of the RNA-binding ability of the related proteins. Therefore, we expanded the RBP2GO database (https://RBP2GO-2-Beta.dkfz.de) to allow the users to browse the information related to RBDs and Rfam IDs in an intuitive manner as well as to use these as search or filter criteria. Therefore, we added a dedicated section which contains a search tool to display the RNA-binding status of InterPro IDs to the sidebar (Figure 8C-D). We also implemented a tab panel in the species-specific “Protein search” to provide the available information pertaining to the InterPro ID annotations as well as the RNA-binding properties of each domain, the presence of disordered regions in the proteins and the RBP2GO composite score (Figure 8E). Protein-specific links to the InterPro and MobiDB databases are provided to simplify access to complementary data if needed. Finally, we generated several options in the “Advanced search” for all the species. The users can navigate through the proteins of the database using a list of specific InterPro IDs and can specifically select proteins that contain RBDs and/or Rfam IDs.

## DISCUSSION

RBPs are important interaction partners for RNA transcripts from the sites of RNA production until their degradation. It is evident that RNA-protein interactions are relevant in numerous key cellular processes. Thus, defects in RBPs are often associated with severe disease phenotypes such as muscular dystrophies, neurological disorders or cancers (3,78). Consequently, it is essential to investigate the composition of ribonucleoprotein complexes in order to understand the molecular mechanisms and functions governed by RNA-protein interactions.

In this study, we present a detailed analysis of RNA-binding features in RBPs from 13 species including datasets generated in both, eukaryotes and prokaryotes. Based on experimental knowledge on RBPs and non-RBPs, we compiled a list of selected RBDs and Rfam IDs. Using the previously defined RBP2GO score (22), we show that the presence of RBDs in proteins or their association with Rfam ID particularly highlights their RNA-binding potential. The number of RBDs per RBP also shows a positive correlation with the RBP2GO score in human RBPs, which is supported by the modularity of canonical RBDs, with several of them often working together within the same protein to more efficiently bind to the target RNAs (26,79).

The distribution of the RBDs between RBPs and non-RBPs in the different species, as well as the higher proportion of non-RBPs with RBD in the less-studied species, confirms the importance of a good study coverage for the proper classification of the proteins into RBPs and non-RBPs. One notable exception to this statement is *Escherichia coli*, which had only 48 non-RBPs with RBD despite a low study coverage with only three RBP screens (16,80,81). This unique situation for *Escherichia coli* may be explained by a significantly lower number of RBD-containing proteins as compared to other species. In the *Escherichia coli* dataset, there are only 247 RBD-containing proteins in total as compared to 1938 proteins in the *Danio rerio* dataset with also three RBP screens. The difference may be due to the size of the respective proteomes (4522 proteins for *Escherichia coli* versus 25801 for *Danio rerio*). However, almost 80% of the RBD-containing proteins in *Escherichia coli* have been identified (Supplementary Table S20), as compared to only 11% in *Dani rerio* (Supplementary Table S4). The high RBP identification rate in *Escherichia coli* seems to be due to one study (80), which detected 1107 RBPs using a method based on silica beads purification. These RBPs cover 98% of the RBD-containing RBPs (185 out of 189) and 78% of the RBD-containing proteins in general (185 out of 237). The same study identified 1448 RBPs in *Saccharomyces cerevisiae*, that cover 73% of the RBD-containing RBPs (438 out of 604) in this species (Supplementary Table S20). The RBPs from this study, that do not contain RBDs, will nonetheless need further experimental validation.

Using the list of selected RBDs, we extended the pool of RBP candidates by identifying the non-RBPs with RBD in all the species of this study. This added 1465 RBP aspirants to the well-studied species and 6273 RBP aspirants to the less-studied species. These were validated both by a GO term enrichment analysis and a literature search. We noticed that the RBP aspirants had a significantly higher RBP2GO score than the non-RBPs lacking RNA-related feature. Non-RBPs, by definition, contribute to their RBP2GO score solely through their interaction partners (22). Hence, RBP aspirants seem to interact more with RBPs than the non-RBPs lacking any RNA-related annotation. This is in agreement with the previous findings reporting that proteins interacting with RBPs have a high probability to be RBPs themselves (19,25). Finally, we observed that the human non-RBPs were significantly less expressed than the RBPs, including the RBP aspirants. A lower expression level can contribute to the difficulty to capture such proteins in large mass spectrometry-based studies, which can explain why some human RBD-containing proteins had not been detected as RBPs yet, despite more than 40 published datasets (22).

Our analysis of IDRs demonstrated that, among RBPs, the RBPs with RBD were enriched in IDRs as compared to the proteins lacking RBDs, which is in agreement with previous studies (74,82,83). However, our data also showed an enrichment in IDRs for non-RBPs with RBD as compared to the RBPs without RBD. Since IDRs confer flexibility to a protein, they can link different domains, such as RBDs as documented for the RNA-binding protein FUS between its RRMs (84). IDRs can also collaborate with canonical RBDs for the recognition of RNA and participate in the specificity of RBPs (85). Based on our current analysis, it seems that IDRs cannot explain the RNA-binding properties of a large number of RBPs lacking RBDs. However, specific types of disordered regions containing LCRs such as arginine/serine (RS) repeats or RGG-boxes can bind to RNA (30,66,86). Such LCRs seem to correlate as well with the presence of RBDs in the proteins. Presently, the MobiDB-lite algorithm, which we used, provides IDR subtypes based on the enrichment in specific amino acids and charge, but not on specific repeats (34,87). Thus, future options to further investigate the impact of IDRs and LCRs in RNA-protein interactions could be to include a detailed analysis of the different repeat subtypes. Altogether, the question of how RBPs without RBD are interacting with RNA remains unclear and of great interest for future investigations, especially for proteins with so far unexpected affinity for RNA and functionally “riboregulated” (67).

Next, we also took advantage of our comprehensive dataset to identify new RBDs within RBPs without RBD but with high RBP2GO scores. Among them, some were found in proteins already known to bind to RNA, such as the histidine kinase/HSP90-like ATPase domain (IPR003594) present in HSP90A (76). The new RBDs were validated using experimental datasets describing RNA-bound protein peptides (13,16,32,45), demonstrating the utility of computational meta-analyses as orthogonal approach besides experimental studies.

Last but not least, we generated the new RBP2GO composite score, which integrates both the previous experimental analysis (components 1 and 2) and the new sequence-based RNA-related analysis (component 3). This new score now allows refining the pool of existing RBPs and identifying new RBPs, which are two independent results. We also applied the RBP2GO composite score to compute a high-confidence list of 2018 RBPs, thereby answering the need for a comprehensive catalog of human RBPs (Supplementary Table S13), which will help selecting candidates for further validation and characterization. In the advanced version of the RBP2GO database, the component of the RBP2GO composite score that is based on the quality factors of the related InterPro IDs was implemented also for species currently lacking an RBP2GO score. We believe that this additional information will facilitate the selection of RBPs for the species with currently low study coverage.

In summary, this study first provides a pan-species list of selected RBDs and Rfam IDs enriched in RBPs and then exploits this resource (i) to refine the list of RBPs from proteome-wide studies with higher confidence, (ii) to predict new RBP aspirants from the group of RBD-containing non-RBPs and (iii) to identify and validate new RBDs from the large group of RBPs lacking selected RBDs. The updated resource is freely available online (https://RBP2GO-2-Beta.dkfz.de) and can be explored in an interactive way. We anticipate that this analysis and the compilation of the information in a unifying platform will stimulate our understanding of RBPs, their functions in key cellular processes and their implications in human diseases.

## DATA AVAILABILITY

RBP2GO 2.0 in a beta version is available at https://RBP2GO-2-Beta.dkfz.de.

The scripts for the main analysis, the production of the figures and the RBP2GO Shiny App are directly available on the RBP2GO-2-Beta database under the DOWNLOAD menu from the sidebar.

## SUPPLEMENTARY DATA

Supplementary data comprises the following items:

**Supplementary Table S1, related to Figure 1**. List of the candidate RNA-related InterPro IDs based on literature and database catalogs.

**Supplementary Table S2, related to Figure 1**. Size of the proteomes of the RBP2GO-listed species.

**Supplementary Table S3, related to Figure 1**. List of selected RBDs.

**Supplementary Table S4, related to Figures 2 to 5**. List of all proteins of the RBP2GO database with information on RBDs, RNA-related family IDs and disorder.

**Supplementary Table S5, related to Supplementary Figure S2**. Expression of human proteins in HeLa cells, downloaded from (https://www.ebi.ac.uk/gxa/experiments/E-PROT-19/Results).

**Supplementary Table S6, related to Supplementary Figure S4.** List of selected RNA-related family IDs (Rfam IDs).

**Supplementary Table S7, related to Figure 6**. Results of the GO term enrichment analysis in the non-RBPs with RBD. If the GO terms are not related to one of the species, the fold-enrichment and the FDR are empty.

**Supplementary Table S8, related to Figure 7**. List of InterPro IDs identified as RBD candidates and their enrichment (three last columns TRUE).

**Supplementary Table S9, related to Figure 7**. List of peptides and domains used for the validation of the new RBDs.

**Supplementary Table S10, related to Figure 8**. Attribution rules of quality factors to RBDs, Rfam IDs, and list of the IDs with their quality factors.

**Supplementary Table S11, related to Figure 8**. Results of the random forest analysis, including the reference lists of proteins used to determine the importance of each parameter.

**Supplementary Table S12, related to Figure S10.** Analysis of the RBP2GO composite score within the groups of RBPs detected using various methods.

**Supplementary Table S13, related to Figure 8**. List of high-confidence human RBPs.

**Supplementary Table S14, related to Figure 8B**. List of the updated proteins of the RBP2GO database with the updated RBDs, RBP2GO score and RBP2GO composite score.

**Supplementary Table S15, related to Figure 2A**. Distribution of RBDs in the RBP2GO species into the three protein categories: RBPs only, non-RBPs only, and both (RBPs and non-RBPs).

**Supplementary Table S16, related to Figure 3 to 5 and Supplementary Figures S3 to S7.** Left (columns A:P): distribution of RBDs and Rfam IDs in the RBP2GO proteins. Right (columns T:AI): distribution of the RBDs and IDRs in the RBP2GO proteins.

**Supplementary Table S17, related to Figure S3.** Analysis of the proteins with RBD and RBP2GO score > 50.

**Supplementary Table S18, related to Figure 4**. Distribution of Rfam IDs in the RBP2GO species into the three protein categories: RBPs only, non-RBPs only, and both (RBPs and non-RBPs).

**Supplementary Table S19, related to Figure 6**. List of non-RBPs with RBD that were validated as RBPs by literature search.

**Supplementary Table S20, related to the Discussion.** List of RBPs detected in the Ec (top table) and Sc (bottom table) proteome-wide studies. The table also include the total number of RBPs detected in Ec and Sc, as well as the total number of RBD-containing proteins (highlighted in yellow).

## AUTHOR CONTRIBUTIONS

M.C.-H. conceived the study and M.C.-H. and S.D. designed and supervised the study and the analysis. E.W., G.K. and M.H. performed the analysis of the RBDs. E.W. performed the analysis of the Rfam IDs, the new RBPs, the new RBDs and their validation and generated the high-confidence list of human RBPs. E.W., G.K. and M.H. produced the final figures, the supplementary tables and implemented the update of the RBP2GO database. E.W. wrote the manuscript, S.D. and M.C.-H. edited the manuscript. All authors have seen and approved the final manuscript.

## ACKNOWLEDGEMENT

The authors sincerely acknowledge the IT core facility of the German Cancer Research Center for their precious support and help in deploying the new version of the RBP2GO database. The authors also deeply thank the lab members, collaborators and users for the feedback on the RBP2GO database, figures and manuscript.

---

**Supplementary Figure S1, related to Figure 1.**
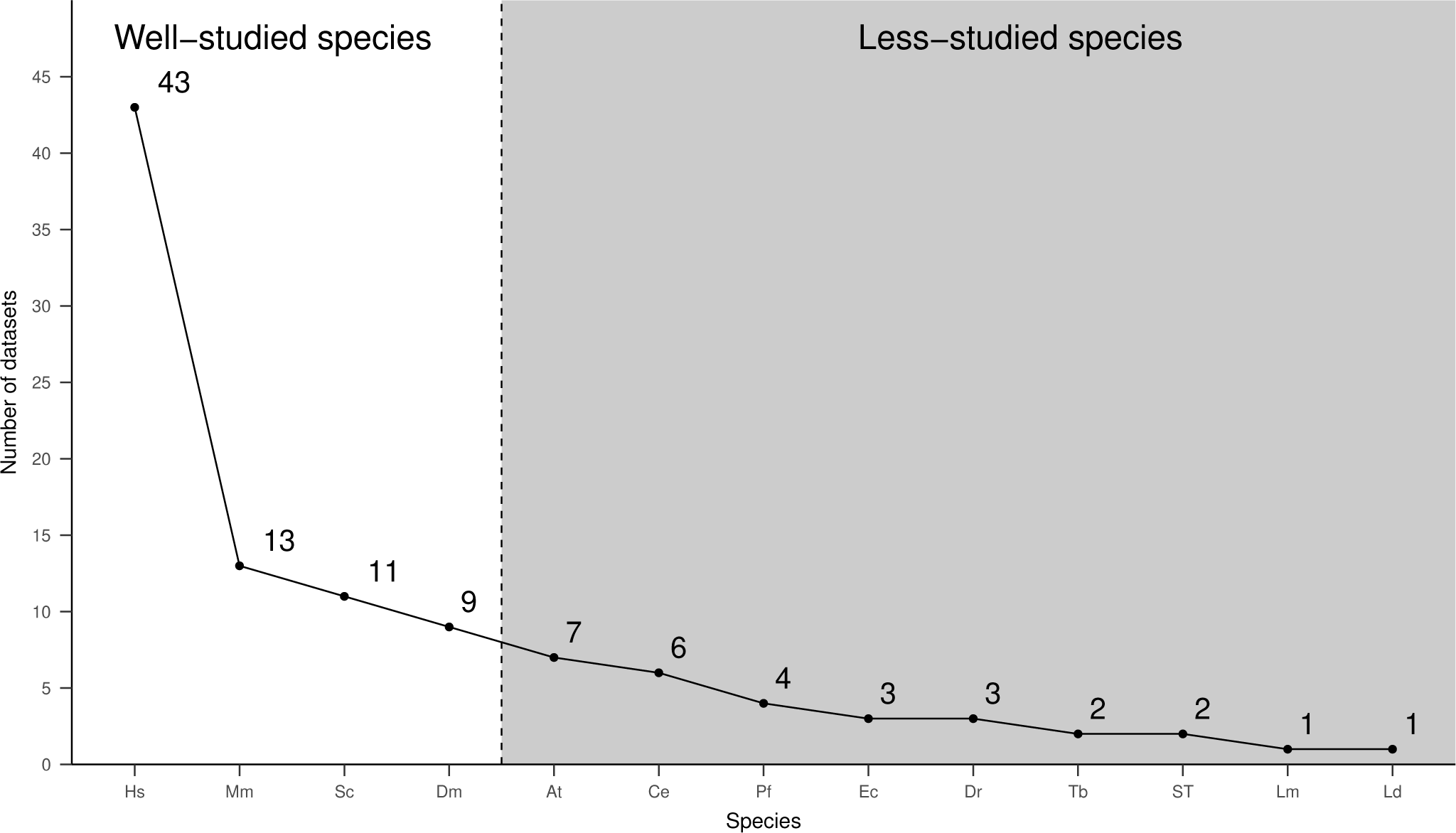
Plot of the number of published proteome-wide RBP screens per species. The vertical dashed line arbitrarily separates the well-studied species (on the left) from the less-studied species (on the right, in the gray area).

**Supplementary Figure S2, related to Figure 2.**
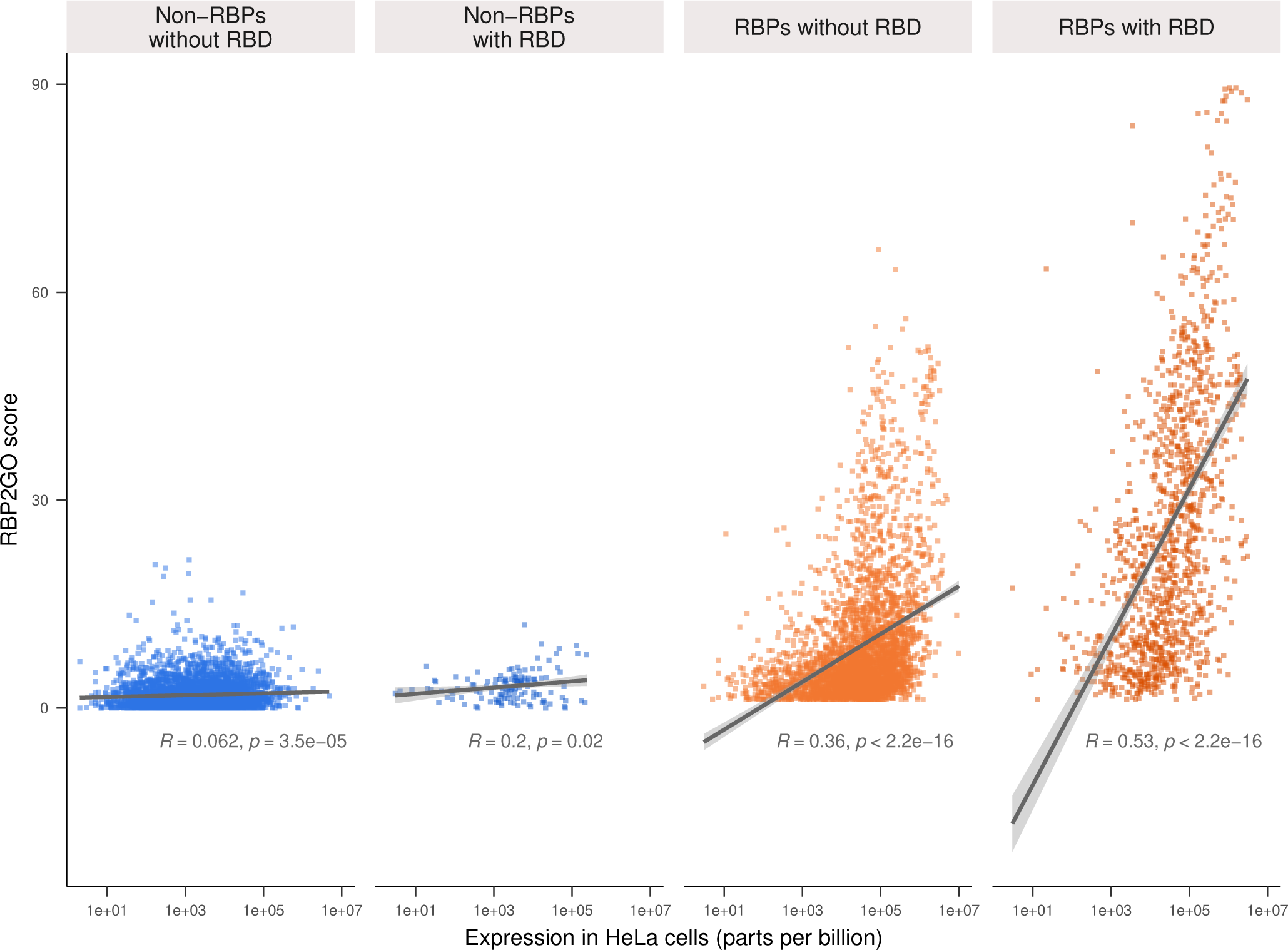
Scatterplot of the RBP2GO scores of human proteins versus their expression in HeLa cells in the four main groups of proteins (non-RBPs without RBD, non-RBPs with RBD, RBPs without RBD and RBPs with RBD). R represents the Pearson’s correlation coefficient. The data used to generate this figure can be found in the Supplementary Table S5.

**Supplementary Figure S3, related to Figure 3.**
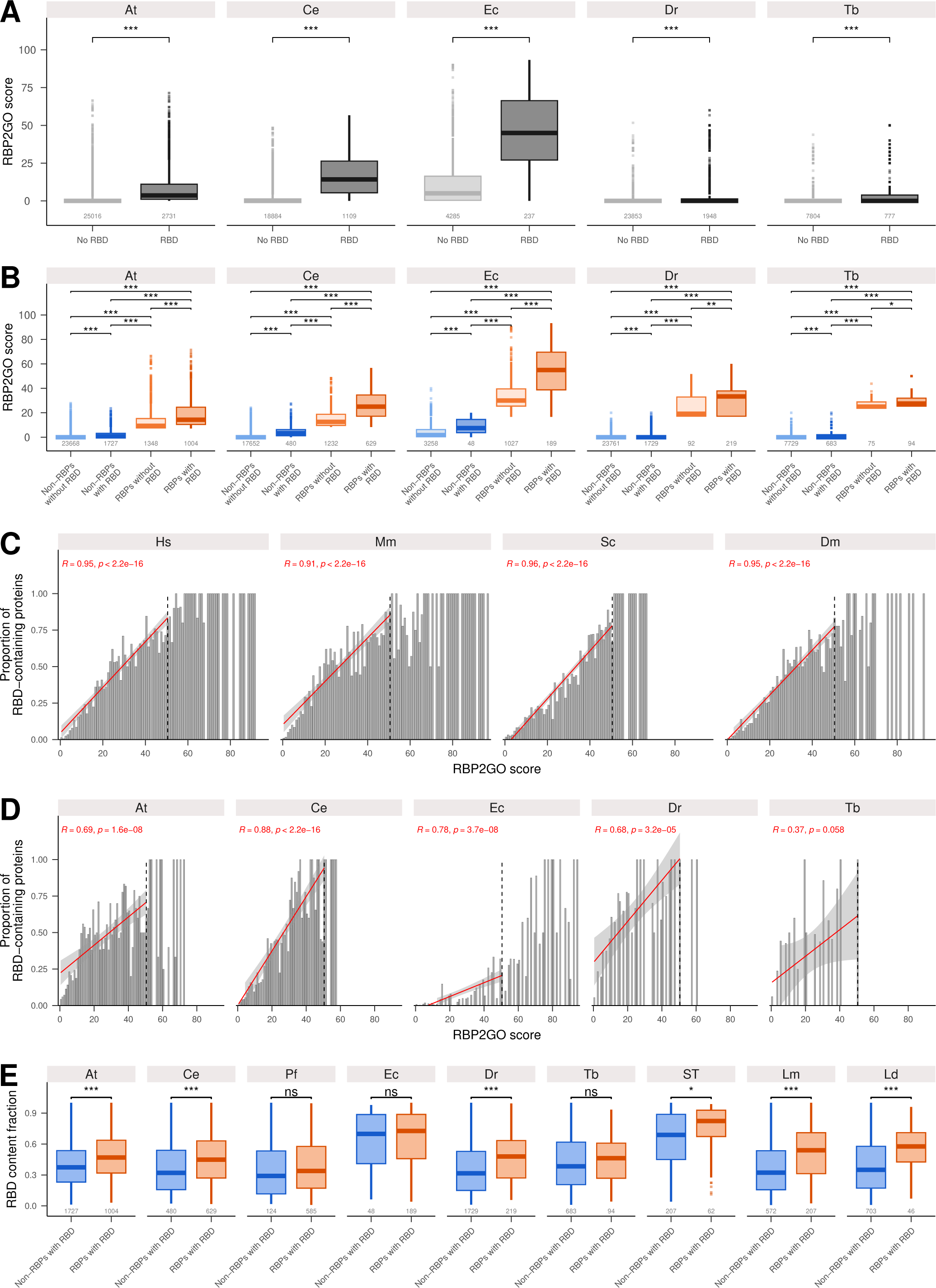
Analysis of the distribution of the RBDs in the proteins of the RBP2GO database species (extended to all species). (**A**) Boxplots representing the distribution of the RBP2GO score in proteins without RBD (light grey) and with RBD (dark grey). (**B**) Same as in (**A**), but separated into non-RBPs without RBD (light blue), non-RBPs with RBD (dark blue), RBPs without RBD (light orange) and RBPs with RBD (dark orange). (**C**) Proportion of RBD-containing proteins for each interval of the RBP2GO score for the well-studied species. The red line represents a linear regression (up to an RBP2GO score of 50). R represents the Pearson’s correlation coefficient. (**D**) Proportion of RBD-containing proteins for each interval of the RBP2GO score for the less-studied species. The red line represents a linear regression (up to a RBP2GO score of 50). R represents the Pearson’s correlation coefficient. (**E**) Boxplots representing the distribution of the number of RBDs per protein in non-RBPs (blue) and RBPs (orange). *, **, *** and **** correspond to *P*-values < 0.05, 0.01, 0.001 and 0.0001, respectively, ns corresponds to non-significant, as resulting from a Wilcoxon rank sum test.

**Supplementary Figure S4, related to Figure 4.**
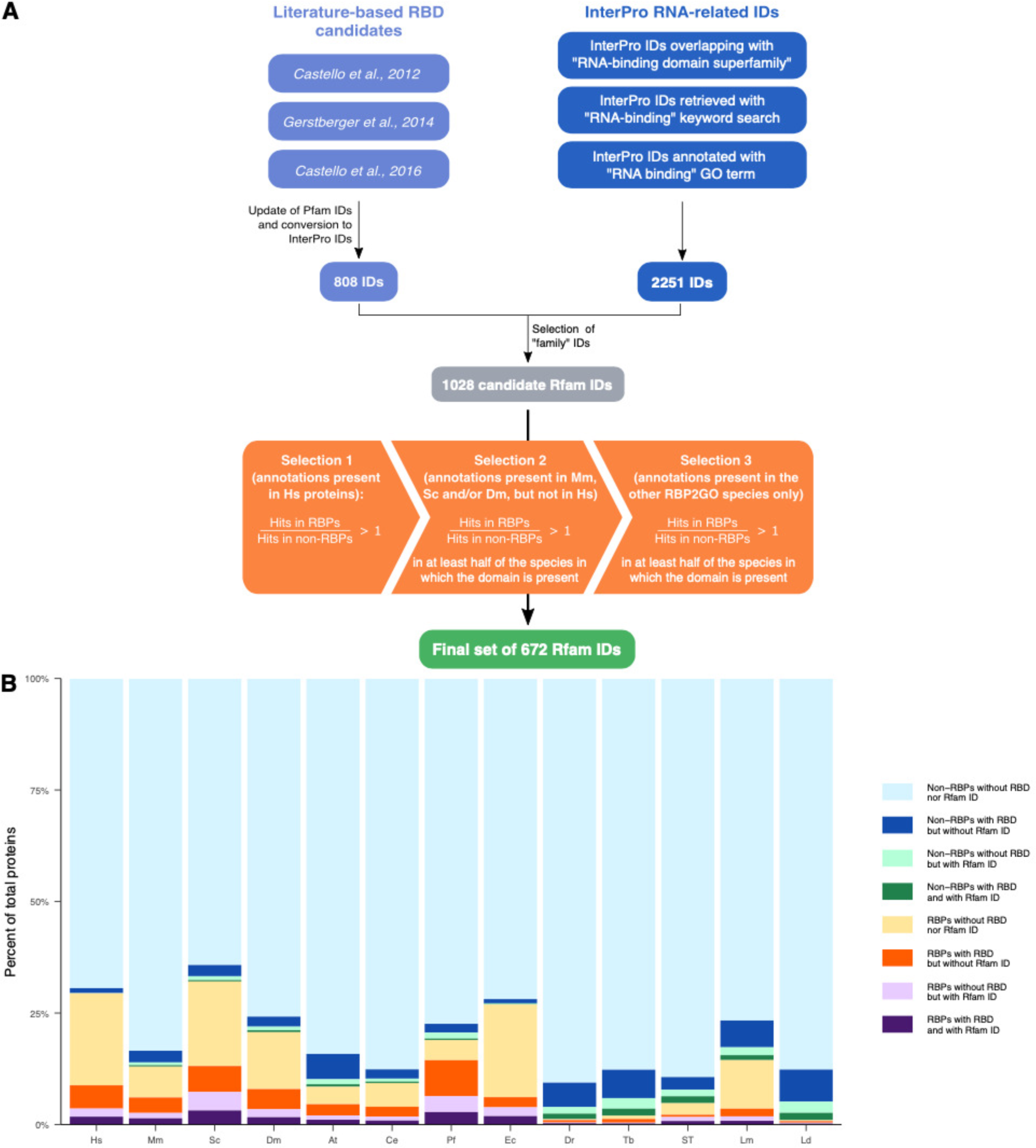
Selection the Rfam IDs. (**A**) Flow chart showing the selection process of the Rfam ID candidates and depicting the three-step selection procedure based on the ratios between RBPs and non-RBPs in the species available in the RBP2GO database (22). The starting lists of Rfam ID candidates (InterPro type = Family) and the final list of selected Rfam IDs are found in Supplementary Tables S1 and S6, respectively. (**B**) Proportion in % of non-RBPs without RBD nor Rfam ID (light blue), non-RBPs with RBD but without Rfam ID (dark blue), non-RBPs without RBD but with Rfam ID (light green), non-RBPs with RBD and Rfam ID (dark green), RBPs without RBD nor Rfam ID (light orange), RBPs with RBD but without Rfam ID (dark orange), RBPs without RBD but with Rfam ID (light purple) and RBPs with RBD and Rfam ID (dark purple) in the different species.

**Supplementary Figure S5, related to Figure 4.**
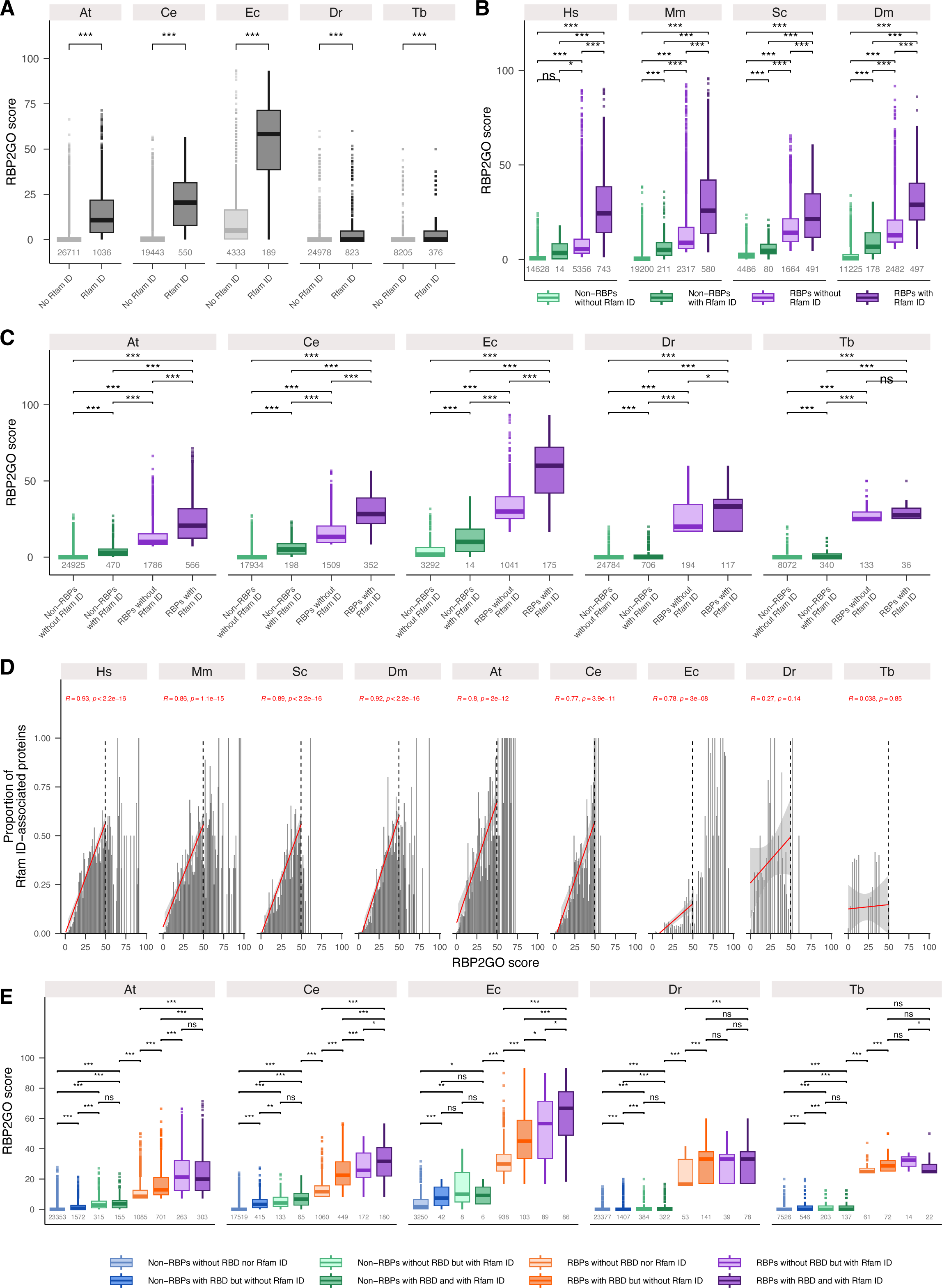
Analysis of the selected Rfam IDs in the proteins of the RBP2GO database species. (**A**) Boxplots representing the distribution of the RBP2GO score in proteins without Rfam ID (no Rfam ID, light grey) and with Rfam ID (Rfam ID, dark grey). (**B**) Boxplots representing the distribution of the RBP2GO score in non-RBPs without Rfam ID (light green), non-RBPs with Rfam ID (dark green), RBPs without Rfam ID (light purple) and RBPs with Rfam ID (dark purple) in the four well-studied species. (**C**) Same as in (**B**) for the less-studied species. (**D**) Proportion of Rfam ID-associated proteins for each interval of the RBP2GO score. The red line represents a linear regression (up to a score of 50). After a score of 50, it does not appear to be linear anymore. R represents the Pearson’s correlation coefficient. (**E**) Boxplots representing the distribution of the RBP2GO score in non-RBPs without RBD nor Rfam ID (light blue), non-RBPs with RBD but without Rfam ID (dark blue), non-RBPs without RBD but with Rfam ID (light green), non-RBPs with RBD and with Rfam ID (dark green), RBPs without RBD nor Rfam ID (light orange), RBPs with RBD but without Rfam ID (dark orange), RBPs without RBD but with Rfam ID (light purple) and RBPs with RBD and with Rfam ID (dark purple). *, **, *** and **** correspond to *P*-values < 0.05, 0.01, 0.001 and 0.0001, respectively, ns corresponds to non-significant, as resulting from a Wilcoxon rank sum test.

**Supplementary Figure S6, related to Figure 5.**
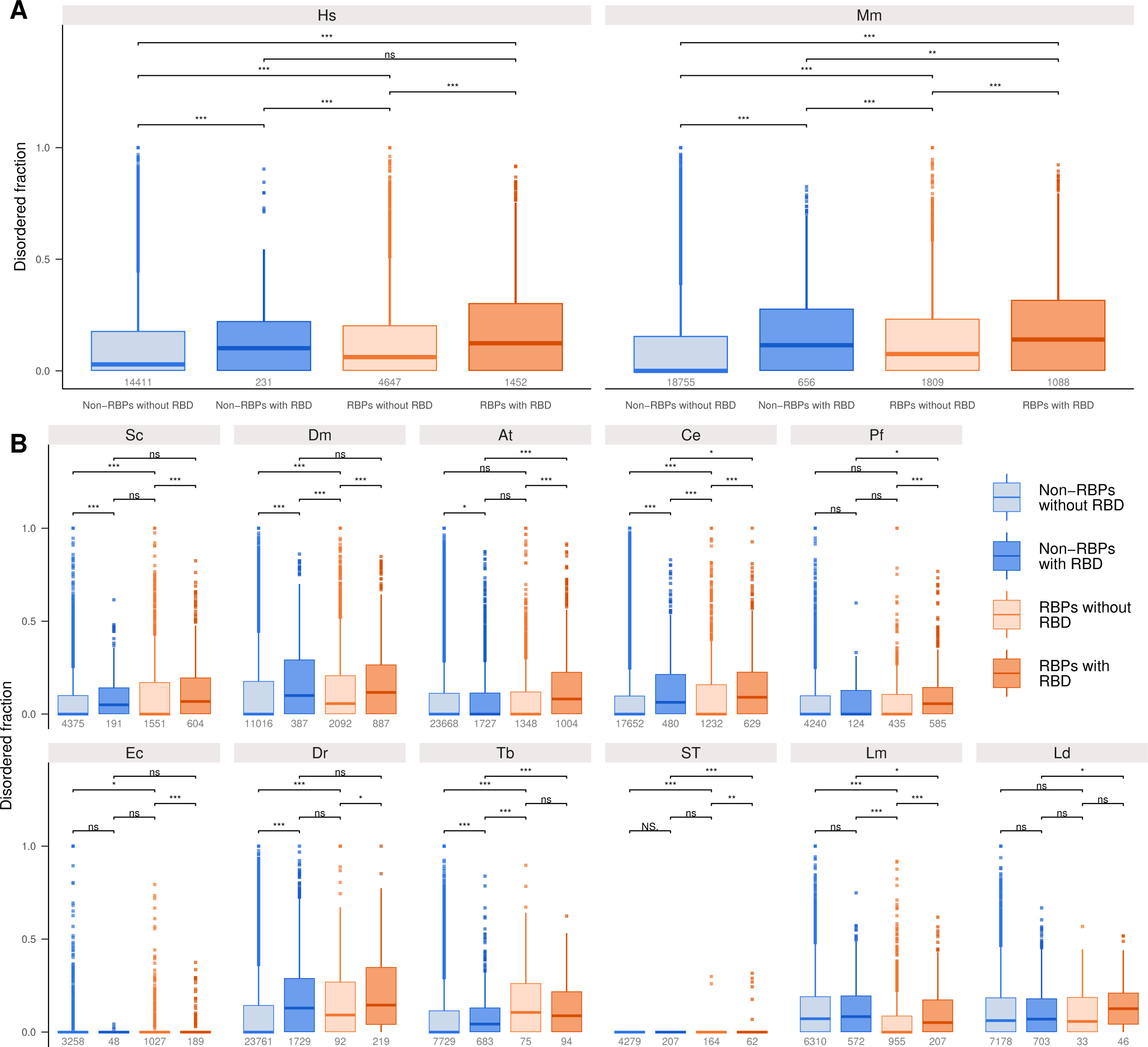
(**A**) Analysis of the disordered fraction in the proteins of the RBP2GO database species (in human and mouse). Boxplots representing the disordered fraction in proteins classified in the following four groups: non-RBPs without RBD (light blue), non-RBPs with RBD (dark blue), RBPs without RBD (light orange) and RBPs with RBD (dark orange). The proteins containing no disordered region were also included. The numbers given in grey are the number of proteins in each group. (**B**) Same as in (**A**) extended to all other species. *, **, *** and **** correspond to *P*-values < 0.05, 0.01, 0.001 and 0.0001, respectively, ns corresponds to non-significant, as resulting from a Wilcoxon rank sum test.

**Supplementary Figure S7, related to Figure 5.**
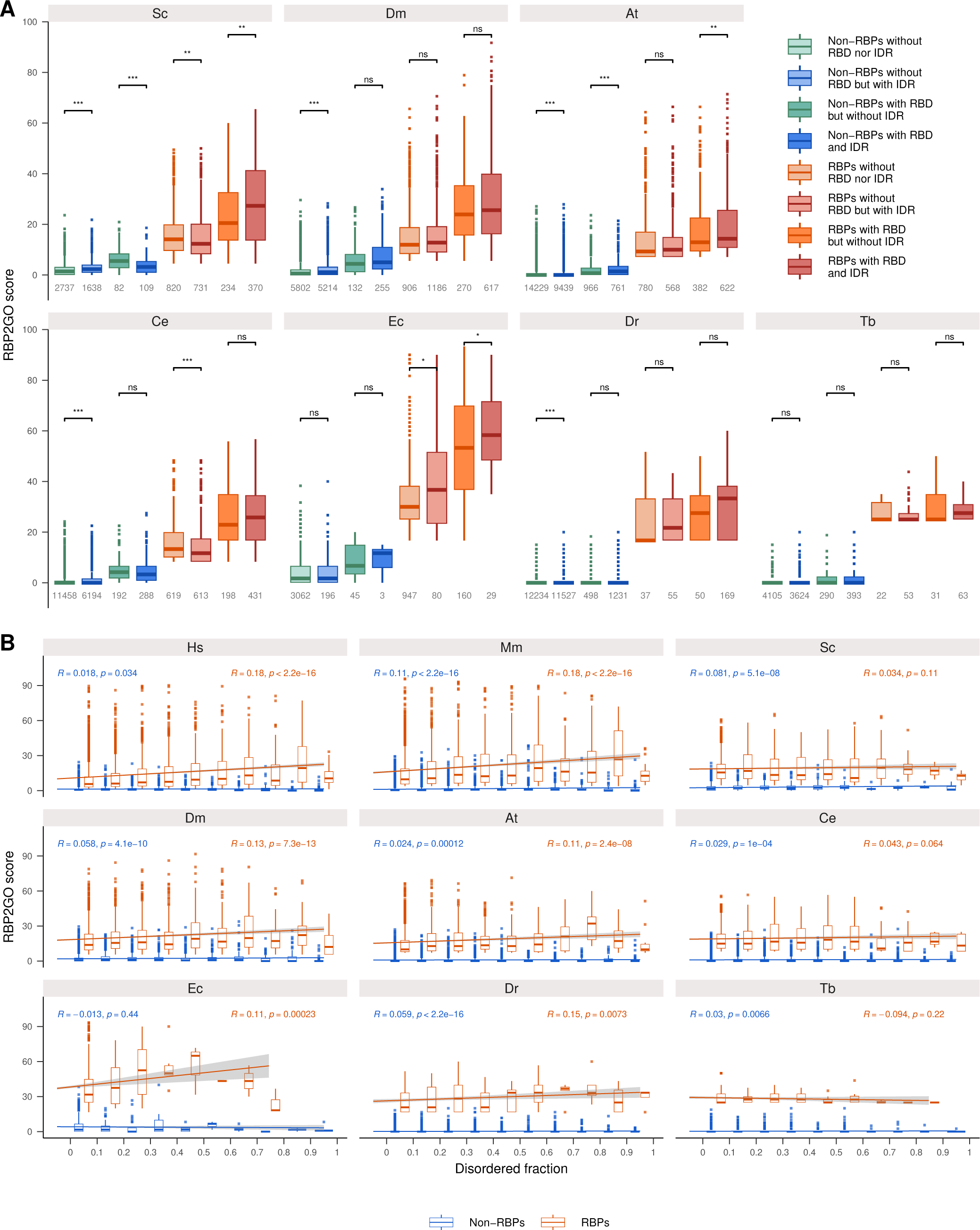
Analysis of the RBP2GO score of the proteins vs the presence of IDR in the proteins of the RBP2GO database species (extended to all species). (**A**) Boxplots representing the distribution of the RBP2GO score in eight groups of proteins: non-RBPs without RBD nor IDR (light green), non-RBPs without RBD but with IDR (light blue), non-RBPs with RBD but without IDR (dark green), non-RBPs with RBD and IDR (dark blue), RBPs without RBD nor IDR (light orange), RBPs without RBD but with IDR (light red), RBPs with RBD but without IDR (dark orange) and RBPs with RBD and IDR (dark red). *, **, *** and **** correspond to *P*-values < 0.05, 0.01, 0.001 and 0.0001, respectively, ns corresponds to non-significant, as resulting from a Wilcoxon rank sum test. (**B**) Boxplots representing the distribution of the RBP2GO score for non-RBPs (blue) and RBPs (orange) in function of the disordered fraction of the proteins. R represents the Pearson’s correlation coefficient.

**Supplementary Figure S8, related to Figure 5.**
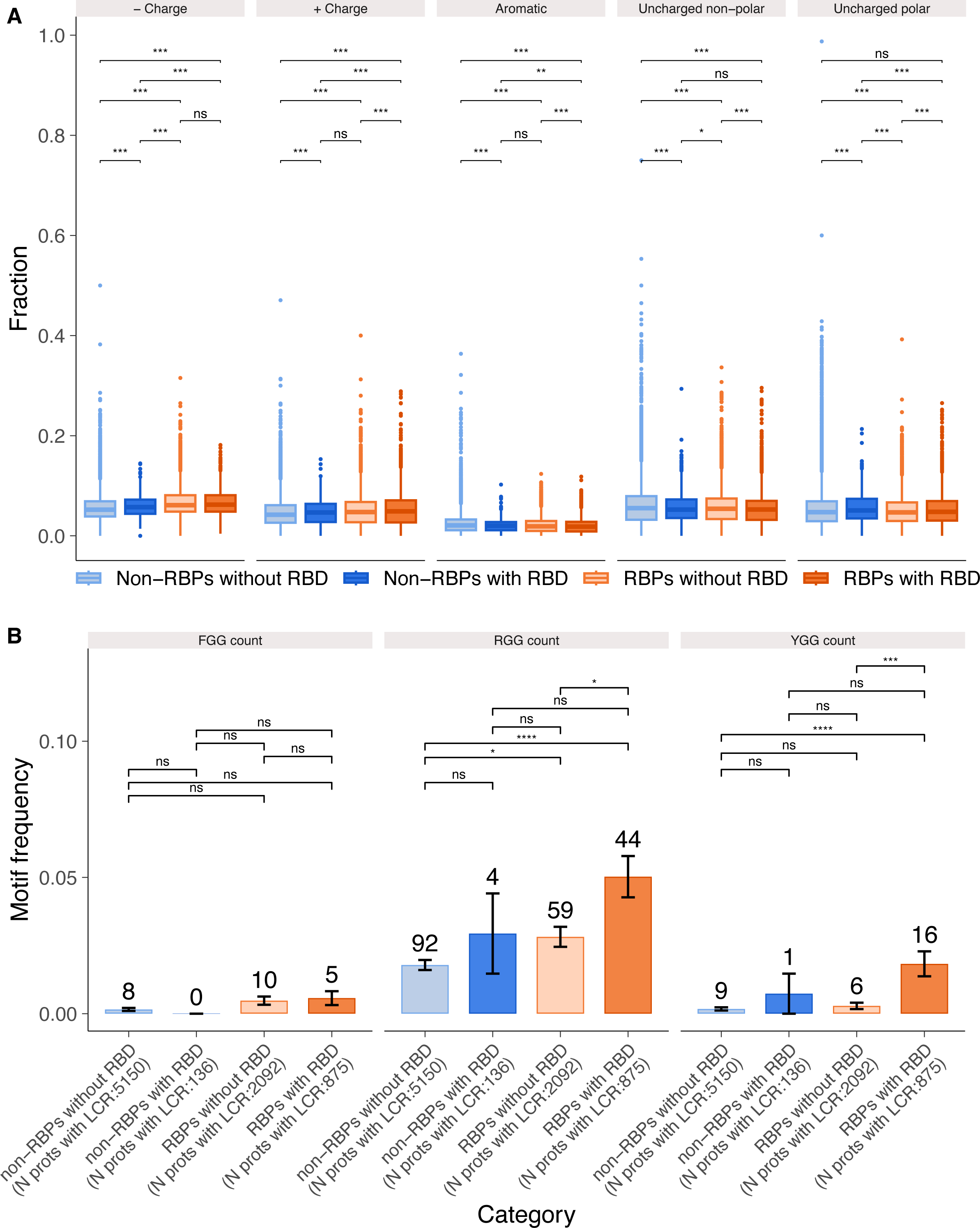
Analysis of the LCR content of the proteins. (**A**) Boxplots showing the distribution of the five groups of amino acids (-charged, + charges, aromatic, uncharged non-polar and uncharged polar) in the four main groups of proteins: non-RBPs without RBD (light blue), non-RBPs with RBD (dark blue), RBPs without RBD (light orange) and RBPs with RBD (dark orange). (**B**) Frequency of the motifs FGG, RGG and YGG in the four main groups of proteins as above in (A). *, **, *** and **** correspond to *P*-values < 0.05, 0.01, 0.001 and 0.0001, respectively, ns corresponds to non-significant, as resulting from a Wilcoxon rank sum test.

**Supplementary Figure S9, related to Figure 8.**
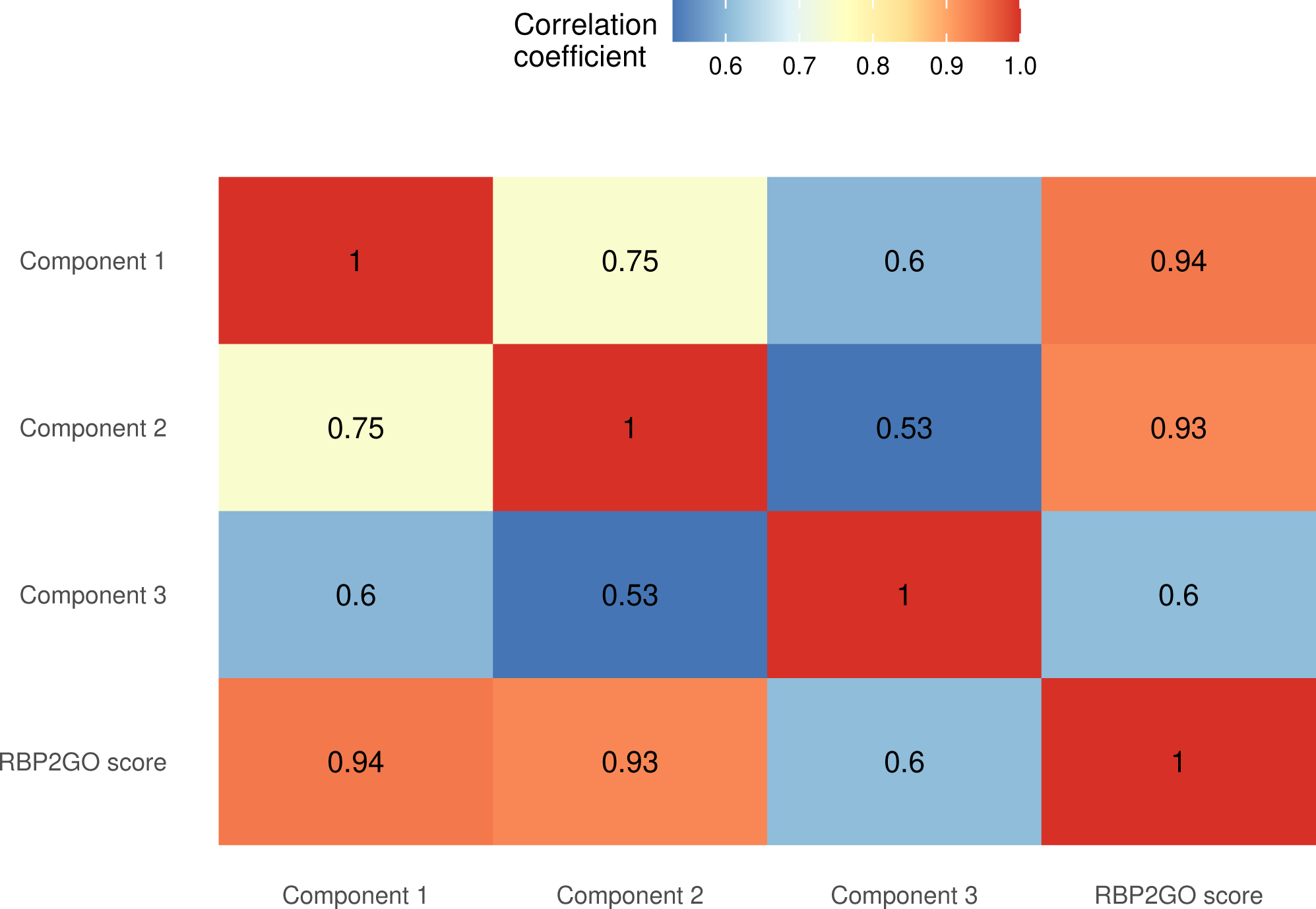
Pearson’s correlation coefficients between the three components of the RBP2GO composite score and the RBP2GO score. The coefficients are depicted from blue (low) to red (high).

**Supplementary Figure S10, related to Figure 8.**
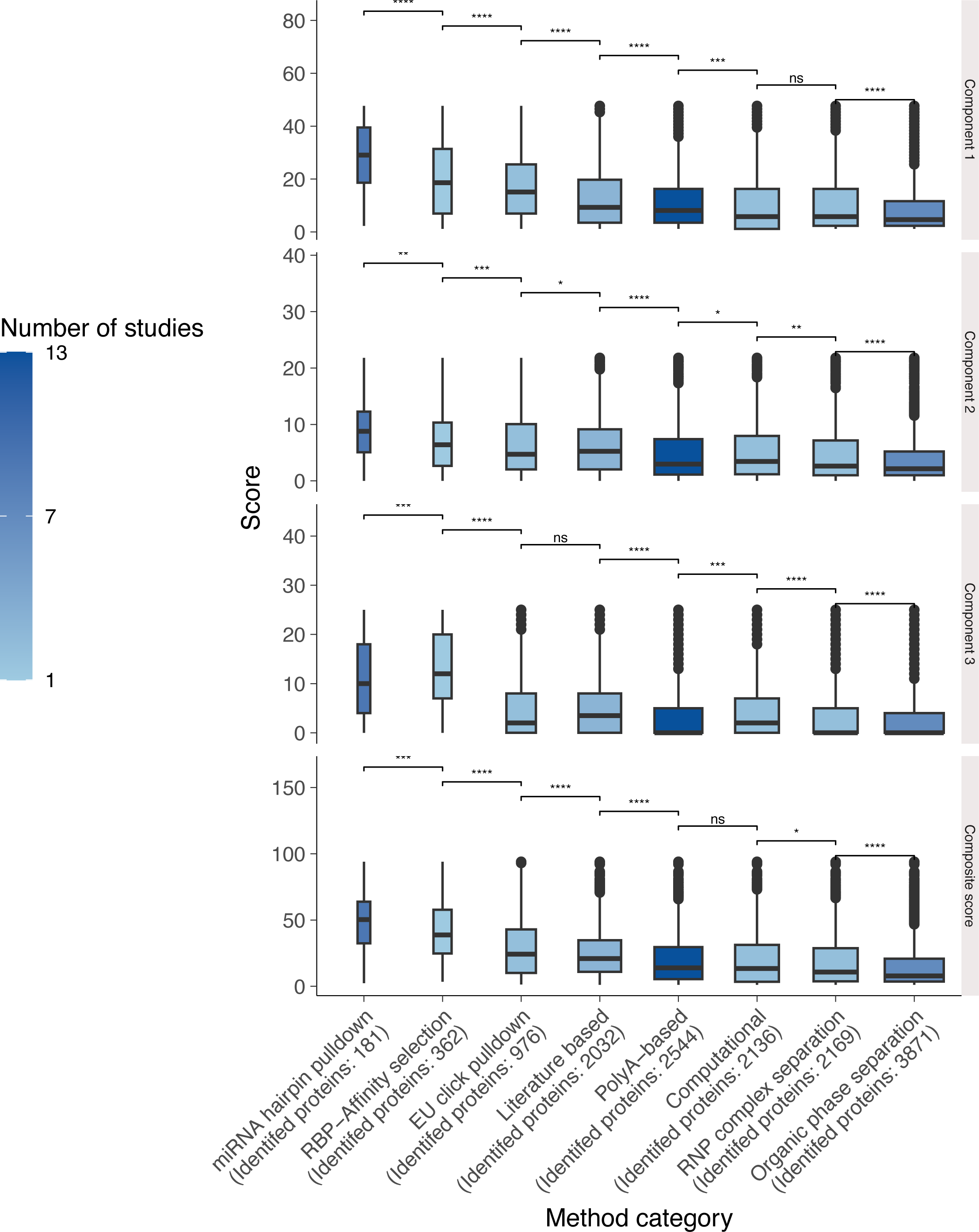
Boxplots representing the distribution of the three components and the RBP2GO composite score in the proteins as detected using various proteome-wide strategies (method category).

**Supplementary Figure S11, related to Figure 8.**
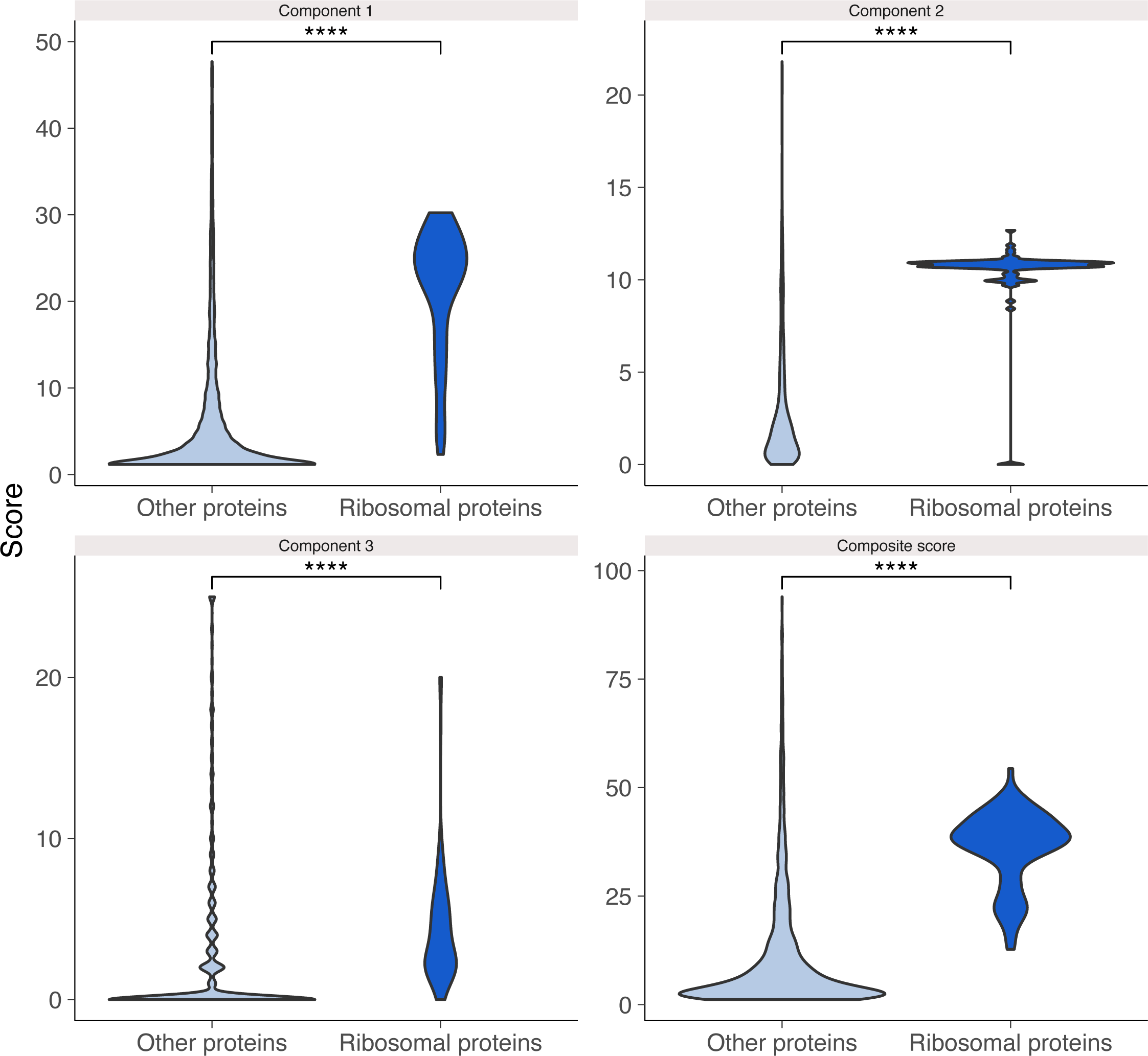
Violin plots showing the distribution of the three components and the RBP2GO composite score in the ribosomal proteins as compared to the other proteins. **** correspond to *P*-values < 0.0001, as resulting from a Wilcoxon rank sum test.

**Supplementary Figure S12, related to Figure 8.**
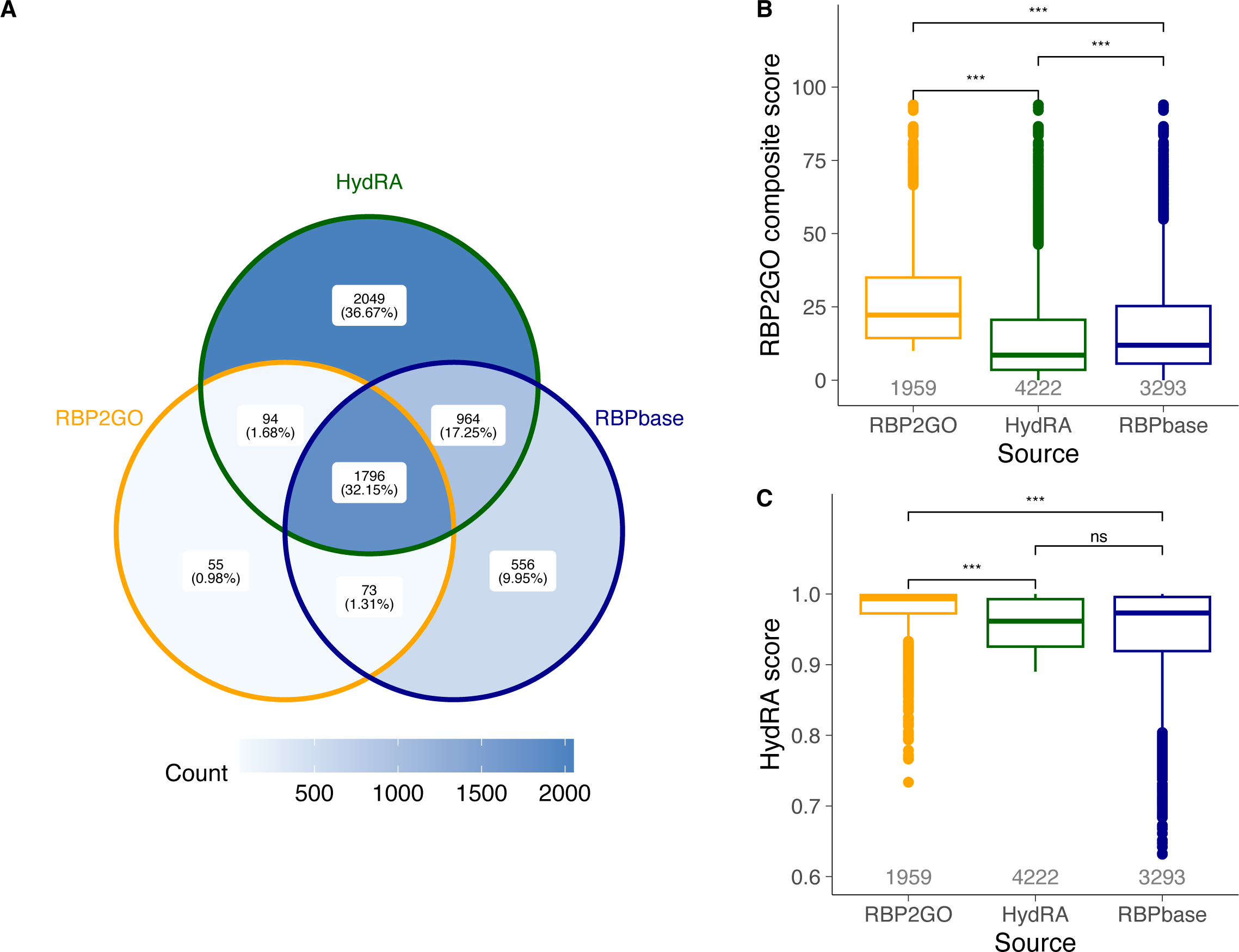
Comparative analysis of RBP2GO, HydRA and RBPbase. (**A**) Venn diagram depicting the overlap between the RBP2GO high-confidence RBPs, the HydRA RBPs and the RBPbase superset of RBPs. (**B**) Boxplot depicting the RBP2GO composite score for these three groups of RBPs (only proteins with available RBP2GO composite score and HydRA score). (**C**) HydRA score for the same three groups of RBPs as in (**B**). *, ** and *** correspond to *P*-values < 0.05, 0.01 and 0.001, respectively, ns corresponds to non-significant, as resulting from a Wilcoxon rank sum test.

